# Dual-sensing genetically encoded fluorescent indicators resolve the spatiotemporal coordination of cytosolic abscisic acid and second messenger dynamics in Arabidopsis

**DOI:** 10.1101/844118

**Authors:** Rainer Waadt, Philipp Köster, Zaida Andrés, Christian Waadt, Gabriele Bradamante, Konstantinos Lampou, Jörg Kudla, Karin Schumacher

## Abstract

Deciphering signal transduction processes is crucial for understanding how plants sense and respond to environmental changes. Various chemical compounds function as central messengers within deeply intertwined signaling networks. How such compounds act in concert remains to be elucidated. We have developed dual-sensing genetically encoded fluorescent indicators (2-In-1-GEFIs) for multiparametric in vivo analyses of the phytohormone abscisic acid (ABA), Ca^2+^, protons (H^+^), chloride (anions), the glutathione redox potential (*E*_GSH_) and hydrogen peroxide (H_2_O_2_). Simultaneous analyses of two signaling compounds in Arabidopsis (*Arabidopsis thaliana*) roots revealed that ABA treatment and uptake did not trigger rapid cytosolic Ca^2+^ or H^+^ fluxes. Glutamate, ATP, Arabidopsis PLANT ELICITOR PEPTIDE (AtPEP1) and glutathione disulfide (GSSG) treatments induced rapid spatiotemporally overlapping cytosolic Ca^2+^, H^+^ and anion fluxes, but except for GSSG only weakly affected the cytosolic redox state. Overall, 2-In-1-GEFIs enable complementary high-resolution in vivo analyses of signaling compound dynamics and facilitate an advanced understanding of the spatiotemporal coordination of signal transduction processes in Arabidopsis.

## INTRODUCTION

Understanding how plants sense and respond to environmental and extracellular fluctuations is key for our strategic progressions to limit the consequences of climate change on plant growth and crop productivity. Plants have evolved complex signal transduction networks that enable the sensing and integration of extracellular signals, and the processing and transduction of the underlying information into physiological-, growth- and developmental responses. Within such signaling networks, spatiotemporal concentration changes of hormones, the divalent cation Ca^2+^ and reactive oxygen species (ROS) mediate various downstream responses (Dodd et al., 2010; Kudla et al., 2010; Shan et al., 2012; Vanstraelen and Benková, 2012; Mittler et al., 2017; Waszczak et al., 2018; Smirnoff and Arnaud, 2019). Among the plant hormones, abscisic acid (ABA) functions as a central regulator of the plant water status (Cutler et al., 2010; Finkelstein et al., 2013; Yoshida et al., 2019). Dynamic concentration changes of signaling compounds require inter- and intracellular transport, including long-distance transport and signaling, that often depend on proton (H^+^) and electrochemical gradients across membranes (Schumacher, 2014; Choi et al., 2016; Sze and Chanroj, 2018). In addition, environmental and cellular H^+^ concentration (pH) can affect plant growth, development and molecular properties (Shavrukov and Hirai, 2016). Therefore, H^+^ may also function in signaling (Sze and Chanroj, 2018).

In plants, hormonal, Ca^2+^, ROS and pH signaling processes are intertwined on several levels (Hauser et al., 2011; Vanstraelen and Benková, 2012; Gilroy et al., 2014; Steinhorst and Kudla, 2014; Edel and Kudla, 2016), for example to regulate stomatal movements or root hair and pollen tube growth (Munemasa et al., 2015; Mangano et al., 2016; Hauser et al., 2017; Michard et al., 2017). In response to the growth hormone auxin, extracellular ATP, touch or wounding, Ca^2+^ signals are accompanied by an apoplastic alkalization and/or cytosolic acidification (Monshausen et al., 2009, 2011; Behera et al., 2018). However, except for the auxin response (Shih et al., 2015; Dindas et al., 2018), the underlying mechanisms are not well understood. Current models suggest that cytosolic Ca^2+^ and extracellular ROS signals are important for cell-to-cell communication and long-distance signaling (Gilroy et al., 2014; Steinhorst and Kudla, 2014). Although some components, such as the ROS producing Arabidopsis (*Arabidopsis thaliana*) RESPIRATORY BURST OXIDASE HOMOLOG D (AtRBOHD), the ion channel TWO PORE CHANNEL1 (TPC1), GLUTAMATE RECEPTOR-LIKE CHANNELS (GLRs), Ca^2+^-DEPENDENT PROTEIN KINASES (CDPKs/CPKs) and CALCINEURIN B-LIKE (CBL) proteins together with CBL-INTERACTING PROTEIN KINASES (CIPKs) have been linked to such processes (Choi et al., 2016; Waszczak et al., 2018), the underlying mechanisms remain unclear.

In order to decipher the coordination and interdependence of signaling processes, it is important to monitor the spatiotemporal dynamics of signaling compounds. Genetically encoded fluorescent indicators (GEFIs) are currently the state-of-the-art technology for high resolution in vivo monitoring of biological processes (Grossmann et al., 2018; Hilleary et al., 2018; Walia et al., 2018). Although the number of GEFIs is steadily increasing, only a fraction has been introduced into plants, and less have been used in simultaneous multiparametric analyses (Okumoto et al., 2012; Kostyuk et al., 2018; Walia et al., 2018). To facilitate the use of GEFIs in multiparametric analyses, we introduce here the concept of dual-sensing genetically encoded fluorescent indicators (2-In-1-GEFIs) that enable the in vivo monitoring of at least two signaling compounds simultaneously. Through the genetic fusion of two distinct indicators, seven 2-In-1-GEFIs were generated that enable time-efficient and complementary in vivo analyses of ABA, Ca^2+^, H^+^, Cl^−^, H_2_O_2_ and the glutathione redox potential (*E*_GSH_) at unpreceded spatiotemporal resolution. Microscopic analyses of these 2-In-1-GEFIs in Arabidopsis roots confirmed their functionality and revealed that extracellularly applied ABA was rapidly taken up, but without discernible effect on cytosolic Ca^2+^ and pH levels. In contrast, treatments with glutamate, ATP, AtPEP1 and GSSG induced spatiotemporally overlapping fluxes of Ca^2+^, H^+^ and Cl^−^, without noticeable rapid effect on the cytosolic redox state.

## RESULTS

### Approaches to optimize FRET-based ABA indicators

Currently available ABA indicators are ABACUS and ABAleon (Jones et al., 2014; Waadt et al., 2014). Because their expression in plants has an impact on ABA signaling and because they exhibit a relatively small signal-to-noise ratio (Waadt et al., 2015), we aimed to optimize these indicators before utilizing them for multiparametric analyses. As testing system, we chose human embryonic kidney (HEK293T) cells that allow for efficient medium throughput plate reader-based screens. Compared to ABAleon2.15 and ABACUS1-2µ used as positive control and non-responsive ABAleon2.15nr as negative control, the initial screening aimed to evaluate deletion variants of ABAleon2.15 and various combinations of five fluorescent protein Förster Resonance Energy Transfer (FRET)-pairs and three sensory domains (SDs; Figure 1A, Supplemental Figures 1A to 1D). 60 min after application of 0 or 100 µM ABA, ABAleon2.15 deletion variants (d1-d3), as well as ABACUS1-2µ and SD2 variants exhibited emission ratio changes even in control conditions. When compared to ABAleon2.15, two indicators (PmTurquoise-SD1-Venus and PmTurquoise-SD1-cpVenus173) exhibited neglectable responses to control treatments, but increased negative emission ratio changes in response to ABA (Figure 1A). Because PmTurquoise-SD1-cpVenus173 (ABAleonSD1-3) differed from ABAleon2.15 only in the sequences that link the sensory domain with the attached fluorescent proteins (Supplemental Figures 1A and 1D), an additional ABAleonSD1-3 linker screening was performed (Figure 1B). This led to the identification of ABAleonSD1-3L21 (linkers LD and T; Supplemental Figure 1E) with similar properties as ABAleonSD1-3 (Figure 1B).

**Figure 1.**
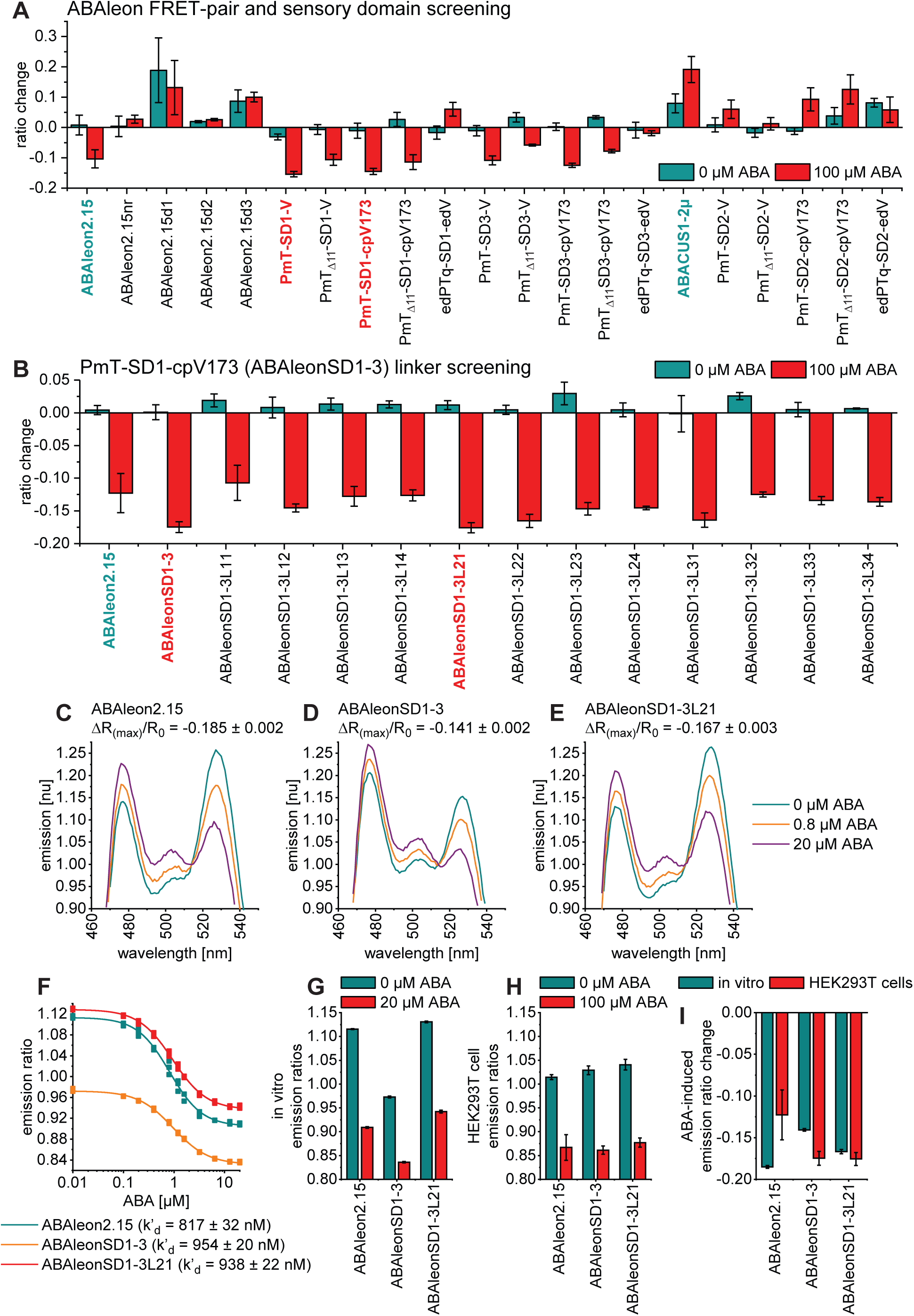
Development of ABAleonSD1-3L21. **(A)** FRET-pair and sensory domain, and **(B)** linker screening of ABA indicator variants after expression in HEK293T cells. Shown are emission ratio changes in response to 60 min treatments with 0 and 100 µM ABA. Reference indicators are shown in cyan and new candidates in red. Information on ABA indicator topologies is given in Supplemental Figure 1. **(C to E)** Representative ABA-dependent normalized in vitro emission spectra of ABAleons with indicated maximum emission ratio change (ΔR_(max)_/R_0_). **(F)** ABA-dependent in vitro emission ratios and apparent ABA affinities (k’_d_) of ABAleons. **(G and H)** Comparison of ABA-dependent ABAleon emission ratios: **(G)** in vitro and **(H)** in HEK293T cells. **(I)** In vitro and HEK293T cell comparison of ABA-induced maximum emission ratio change. All data are shown as mean ± SD, n = 3.

To corroborate these findings, ABAleon2.15, ABAleonSD1-3 and ABAleonSD1-3L21 proteins were purified from *Escherichia coli* (*E. coli*) and characterized in vitro. Although all three ABAleons were functional (Figures 1C to 1G), their properties in vitro were markedly different from results obtained in HEK293T cells (Figures 1G to 1I). Basal emission ratios (at 0 µM ABA) were not the same in both assay systems (Figures 1G to 1H). ABAleon2.15 exhibited a higher ABA-induced emission ratio change in vitro, while ABAleonSD1-3 responses were larger in the HEK293T cell system (Figure 1I). ABAleonSD1-3L21 responded similar in vitro and in HEK293T cells (Figure 1I). In vitro calibrations revealed that ABAleonSD1-3 responded much weaker to ABA (maximum emission ratio change ΔR_(max)_/R_0_ = −0.141) compared to ABAleon2.15 and ABAleonSD1-3L21 (ΔR_(max)_/R_0_ = −0.185 and −0.167) (Figures 1C to 1E). Apparent ABA affinities of ABAleonSD1-3 (954 nM) and ABAleonSD1-3L21 (938 nM) were slightly lower compared to ABAleon2.15 (817 nM) (Figure 1F). Altogether, the ABA indicator screening led to the identification of ABAleonSD1-3L21. Its advanced properties in HEK293T cells requires therefore further validation in planta.

### Evaluation of FRET-based ABA indicators in Arabidopsis

5-day-old Arabidopsis seedlings expressing ABAleon2.15, ABAleonSD1-3L21 or ABACUS1-2µ in the cytosol and nucleus were compared for their ABA responses in roots. Therefore, spatiotemporal vertical response profiles of emission ratios (R; Figure 2 top) or emission ratio changes normalized to 4 min average baseline recordings (ΔR/R_0_; Figure 2 middle) and overall emission ratio changes (Figure 2 bottom) were acquired in response to 10 µM ABA treatments. ABAleon2.15 and ABAleonSD1-3L21 responded similar to ABA with a sigmoidal emission ratio decrease at a half response time of t_1/2_ ∼ 15 min (Figures 2A and 2B). Note that the calyptra exhibited a much weaker response to ABA compared to the other tissues, likely because of the high cytosolic ABA concentration ([ABA]_cyt_) there (depicted in dark blue; Figures 2A top and 2B top). The root elongation zone however, exhibited the lowest [ABA]_cyt_ (depicted in white). ABACUS1-2µ did not resolve this ABA concentration gradient, and responded slower (t_1/2_ ∼ 29 min), but with larger increasing emission ratio changes that were more pronounced in the meristematic- and early elongation zone (Figure 2C, Supplemental Movie 1). None of the indicators exhibited emission ratio changes in response to control treatments (Supplemental Figure 2). Altogether, we conclude that ABAleons are more suitable for ABA analyses in tissues with low [ABA]_cyt_, whereas ABACUS1-2µ should be preferably used in tissues with high [ABA]_cyt_. Because both ABAleons exhibited similar ABA response patterns, we decided to employ the latest version ABAleonSD1-3L21 for our multiparameter imaging approach.

**Figure 2.**
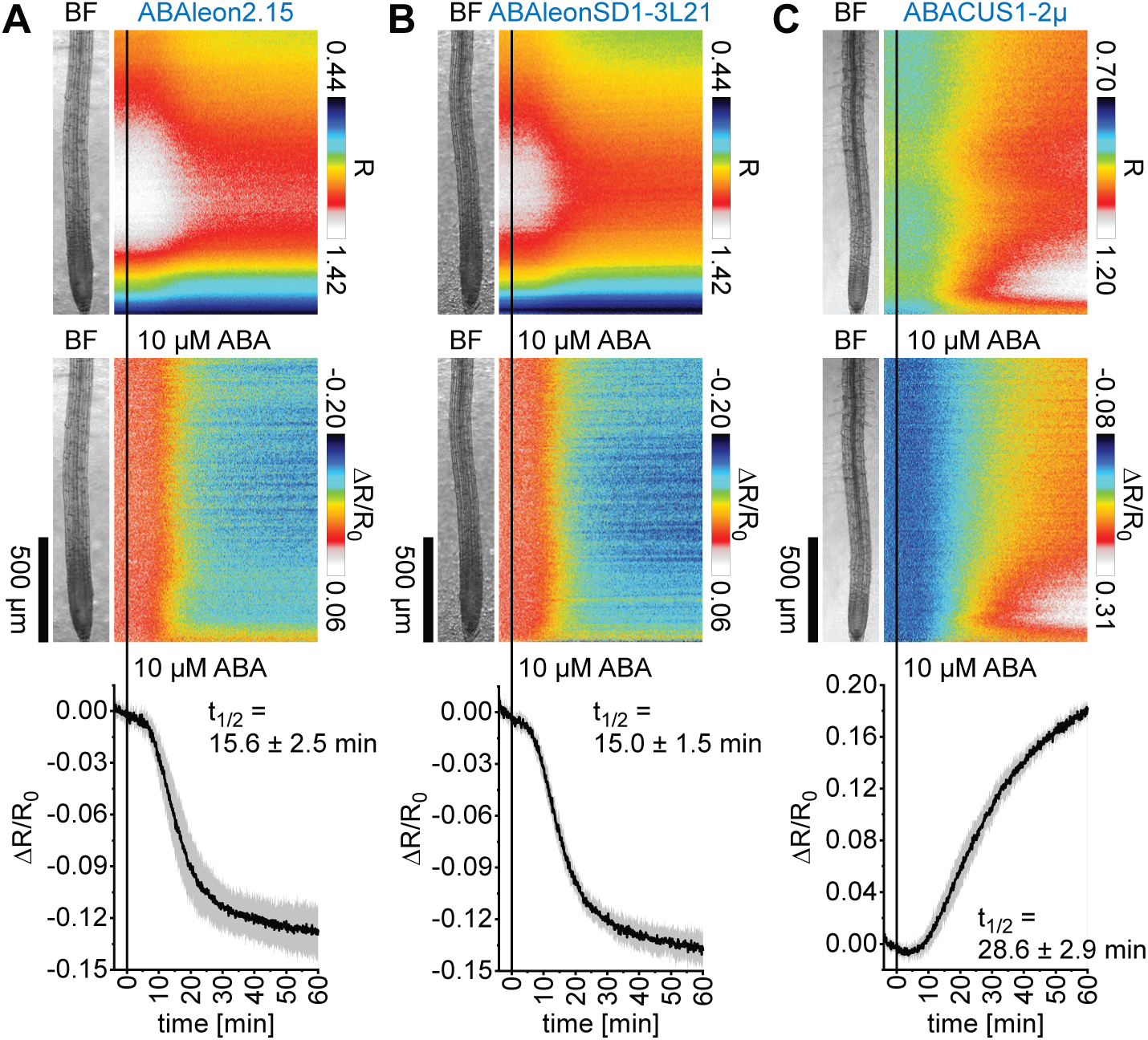
Comparison of ABA indicator ABA responses in Arabidopsis. Five-day-old roots of Arabidopsis expressing **(A)** ABAleon2.15 (n = 9), **(B)** ABAleonSD1-3L21 (n = 8) and **(C)** ABACUS1-2µ (n = 6) were imaged for 64 min at a frame rate of 10 min^-1^ and treated with 10 µM ABA at t = 0 min. Shown are average vertical response profiles of (top) emission ratios (R) and (middle) emission ratio changes (ΔR/R_0_) normalized to 4 min average baseline recordings. An adjacent representative bright field (BF) root image is shown for orientation. (bottom) Full image average emission ratio changes (mean ± SD) with indicated half response times (t_1/2_). A representative experiment is provided as Supplemental Movie 1. Data of 0 µM ABA control experiments are shown in Supplemental Figure 2.

### Concept and design of dual-sensing genetically encoded fluorescent indicators (2-In-1-GEFIs)

Multiparametric analyses of signaling compounds requires the generation of transgenic plants that express several GEFIs simultaneously. Because the generation of transgenic plants is time-consuming and the insertion of multiple transgenes into the Arabidopsis genome often results in epigenetic silencing effects, we aimed to express two GEFIs from one mRNA. Therefore, GEFIs were genetically fused via sequences encoding for a 14 amino acid ASGGSGGTSGGGGS-linker (GSL), or the self-cleaving 22 amino acid P2A linker that enables the expression of two separate polypeptides (Kim et al., 2011). Six GEFIs: ABAleonSD1-3L21 (ABA), R-GECO1 (Ca^2+^; Zhao et al., 2011), Arabidopsis codon optimized red-fluorescing (P)A-17 (H^+^; Shen et al., 2014), E^2^GFP (H^+^ and Cl^−^; Bizzarri et al., 2006), Grx1-roGFP2 (*E*_GSH_; Gutscher et al., 2008) and roGFP2-Orp1 (H_2_O_2_; Gutscher et al., 2009) were combined for the generation of seven 2-In-1-GEFIs: ABAleonSD1-3L21-P2A-R-GECO1 (ABA and Ca^2+^), PA-17-P2A-ABAleonSD1-3L21 (H^+^ and ABA), R-GECO1-GSL-E^2^GFP (Ca^2+^, H^+^ and Cl^−^), PA-17-P2A-Grx1-roGFP2 (H^+^ and *E*_GSH_), PA-17-P2A-roGFP2-Orp1 (H^+^ and H_2_O_2_), R-GECO1-P2A-Grx1-roGFP2 (Ca^2+^ and *E*_GSH_) and R-GECO1-P2A-roGFP2-Orp1 (Ca^2+^ and H_2_O_2_). See Supplemental Data Sets 1B and 1C for information about constructs and transgenic Arabidopsis plants. In the following, we will describe the application of these 2-In-1-GEFIs in Arabidopsis and highlight the resulting biological findings.

### ABA does not trigger rapid Ca^2+^ or pH changes in roots

To test the functionality of the 2-In-1-GEFIs, we first studied the interrelation of ABA with Ca^2+^ in roots. Therefore, ABAleonSD1-3L21-P2A-R-GECO1 seedlings were monitored in response to 10 µM ABA, which induced a typical ABAleonSD1-3L21 emission ratio decrease (Figure 3A left). The Ca^2+^ indicator R-GECO1 did not respond to this treatment (Figure 3A right). However, subsequent 1 µM indole-3-acetic-acid (IAA; auxin) treatment at 30 min induced a biphasic Ca^2+^ signal that initiated in the root elongation zone and spread from there to neighboring regions (Figure 3A right, Supplemental Movie 2), as observed before (Waadt et al., 2017).

**Figure 3.**
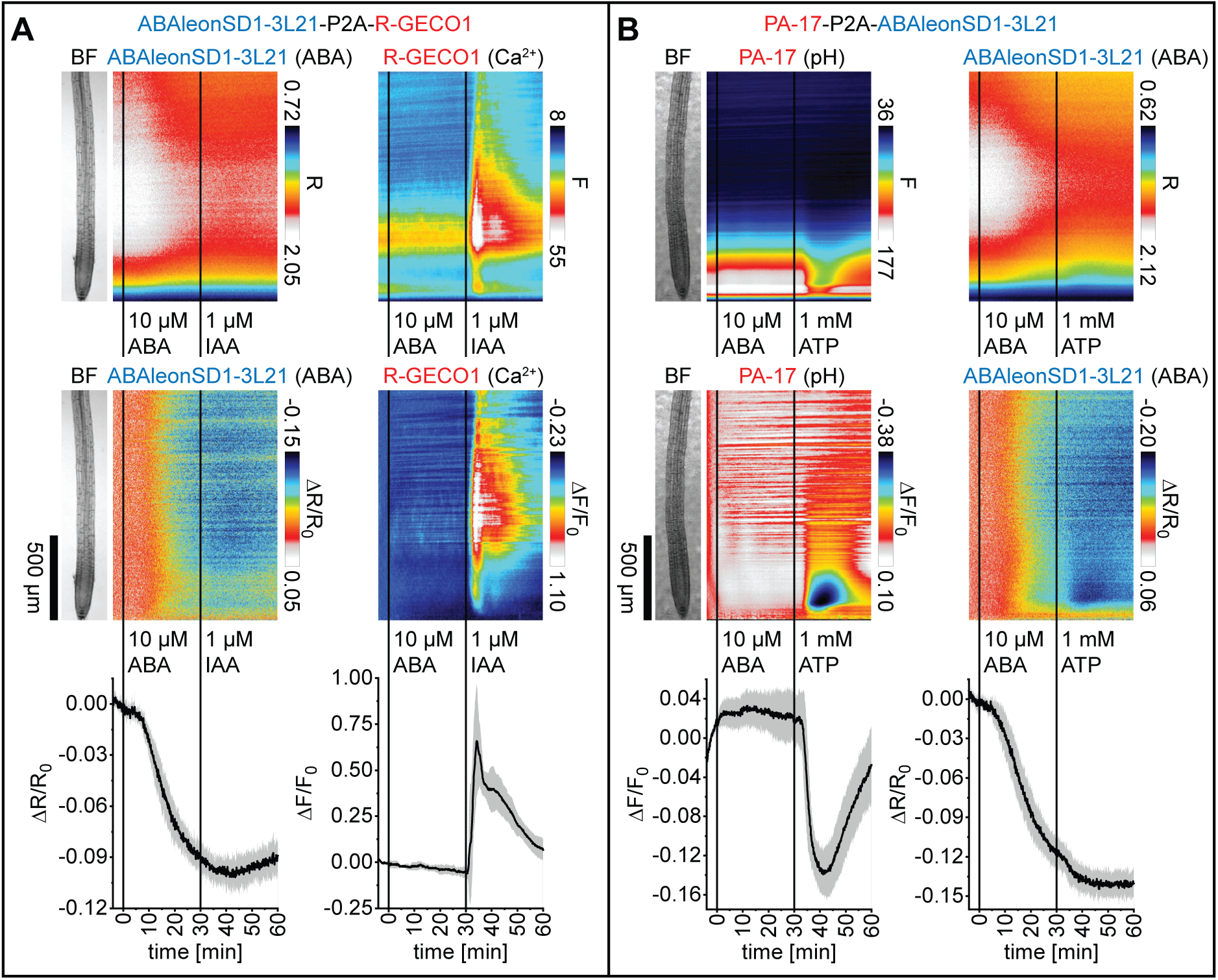
Application of ABA does not trigger rapid changes of cytosolic Ca^2+^ or pH in Arabidopsis roots. Analyses of five-day-old roots of Arabidopsis expressing **(A)** ABAleonSD1-3L21-P2A-R-GECO1 (ABA and Ca^2+^; n = 5) in response to 10 µM ABA (t = 0 min) and 1 µM IAA (t = 30 min), and **(B)** PA-17-P2A-ABAleonSD1-3L21 (pH and ABA; n = 8) in response to 10 µM ABA (t = 0 min) and 1 mM ATP (t = 30 min). Images were acquired for 64 min at a frame rate of 10 min^-1^. Shown are average vertical response profiles of (top) emission ratios (R) or fluorescence emissions (F) and (middle) signal changes (ΔR/R_0_ or ΔF/F_0_) normalized to 4 min average baseline recordings. An adjacent representative bright field (BF) root image is shown for orientation. (bottom) Full image signal changes (mean ± SD). Representative experiments are provided as Supplemental Movies 2 and 3.

To investigate the effect of ABA on cytosolic pH, PA-17-P2A-ABAleonSD1-3L21 seedlings were treated with 10 µM ABA. From 0-30 min after ABA treatment PA-17 fluorescence remained unchanged (Figure 3B left). However, in response to 1 mM ATP PA-17 fluorescence emission decreased, indicating a rapid and transient cytosolic acidification with a maximum pH drop in the root meristematic zone that also spread to the elongation zone (Figure 3B left). In this experiment, ABAleonSD1-3L21 exhibited a typical ABA response pattern that was not affected by ATP (Figure 3B right, Supplemental Movie 3). These experiments established the functionality of both 2-In-1-GEFIs and revealed that ABA does not trigger rapid cytosolic Ca^2+^ or pH changes in roots.

### Auxin, ATP and glutamate treatments trigger spatiotemporally overlapping fluxes of Ca^2+^, H^+^ and Cl^−^

Next, we used R-GECO1-GSL-E^2^GFP to simultaneously monitor Ca^2+^, H^+^ and Cl^−^ fluxes. Because anions, such as Cl^−^, quench the fluorescence of E^2^GFP and because its excitation ratiometric pH readout is Cl^−^ independent, E^2^GFP provides a means to simultaneously assess cytosolic H^+^ and Cl^−^ changes (Bizzarri et al., 2006; Arosio et al., 2010). In response to 1 µM IAA, R-GECO1 reported biphasic Ca^2+^ signals in the root elongation zone that travelled to neighboring regions. Subsequent 1 mM ATP treatments triggered Ca^2+^ signals in the calyptra and meristematic zone that proceeded shootward (Figure 4A left). Interestingly, both Ca^2+^ signals coincided with a cytosolic acidification reported by E^2^GFP (Figure 4A middle, Supplemental Movie 4). IAA also induced a Cl^−^ influx, indicated by a E^2^GFP fluorescence emission decrease in the entire imaged root, with a maximum decrease in the meristematic zone (Figure 4A right). ATP however induced Cl^−^ influx in the upper elongation zone and above, but Cl^−^ efflux in the lower elongation- and meristematic zone (Figure 4A right). Correlation analyses of the initial 15 min during the IAA response indicated a remarkable spatiotemporal overlap of Ca^2+^, H^+^ and Cl^−^ influx in the meristematic and elongation zone (Figure 4B).

**Figure 4.**
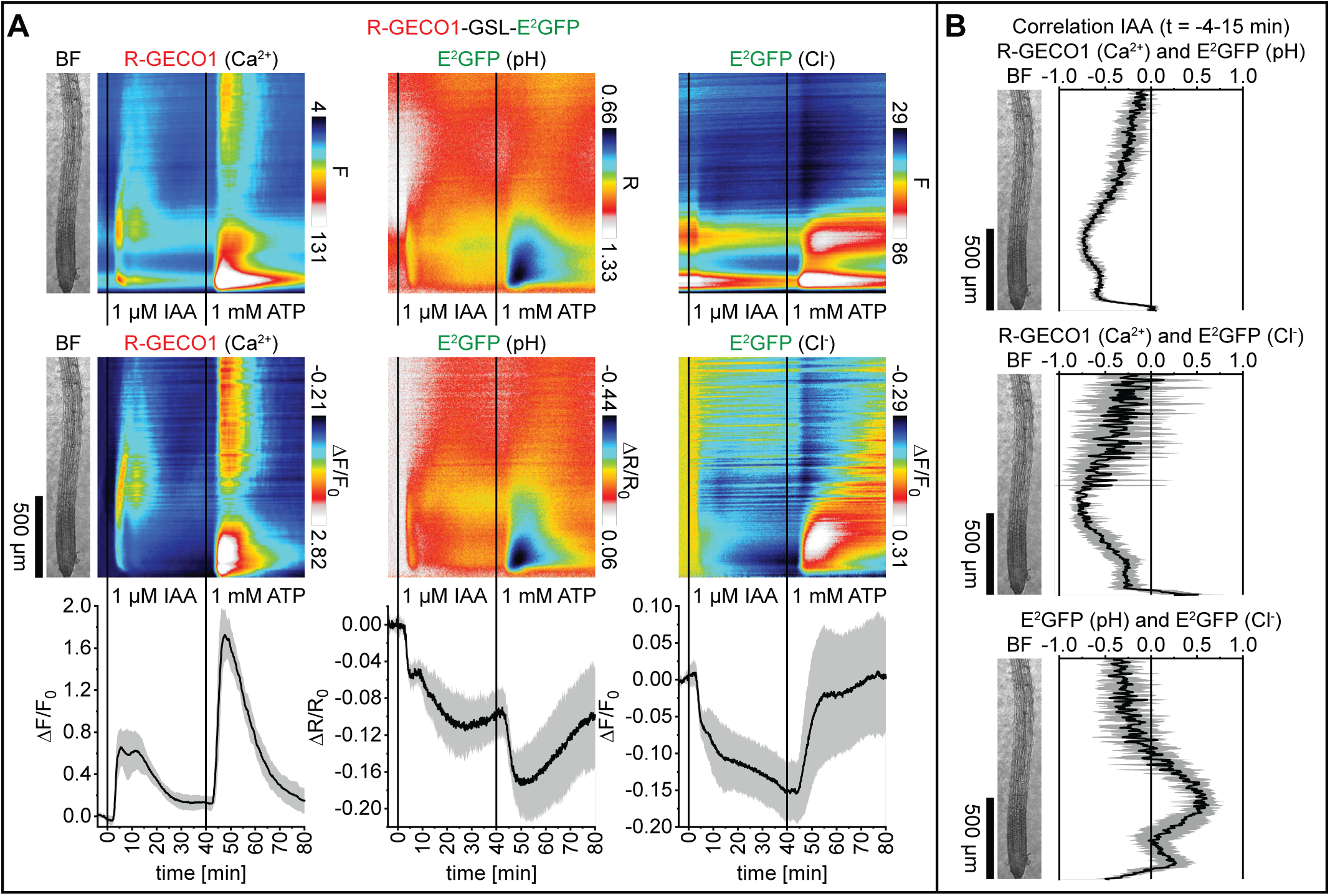
Application of auxin and ATP triggers cytosolic Ca^2+^, H^+^ and Cl^−^ fluxes. Analyses of five-day-old roots of Arabidopsis expressing R-GECO1-GSL-E^2^GFP (Ca^2+^, pH and Cl^−^) in response to 1 µM IAA (t = 0 min) and 1 mM ATP (t = 40 min; n = 6). Images were acquired for 84 min at a frame rate of 10 min^-1^. **(A)** Average vertical response profiles of (top) fluorescence emissions (F) or emission ratios (R), (middle) signal changes (ΔF/F_0_ or ΔR/R_0_) normalized to 4 min average baseline recordings, and (bottom) full image signal changes (mean ± SD). **(B)** Spatiotemporal Pearson correlation analyses (mean ± SD) of indicated GEFI responses during the application of IAA (t = −4-15 min). An adjacent representative bright field (BF) root image is shown for orientation. A representative experiment is provided as Supplemental Movie 4.

In additional experiments, the effect of 1 mM glutamate was assessed. Compared to IAA, glutamate treatments triggered a more expanded and rapid Ca^2+^ transient that arrived in a wave-like shape from upper root regions (Figure 5A left, Supplemental Movie 5). H^+^ and Cl^−^ also displayed a rapid and transient initial influx with a maximum acidification in the meristematic zone, followed by a weak transient alkalization in the early elongation zone and a prolonged H^+^ and Cl^−^ influx (Figure 5A left, Supplemental Movie 5). During the initial 10 min of the glutamate response, Ca^2+^ and H^+^ influx exhibited a noticeable spatiotemporal overlap/correlation in the meristematic- and early elongation zone (Figure 5B left). Subsequent responses to ATP, used as positive control, were as observed before (Figures 4A and 5A). Correlation analyses indicated a remarkable coincidence of Ca^2+^ and H^+^ influx and Cl^−^ efflux in the meristematic zone (Figure 5B right).

**Figure 5.**
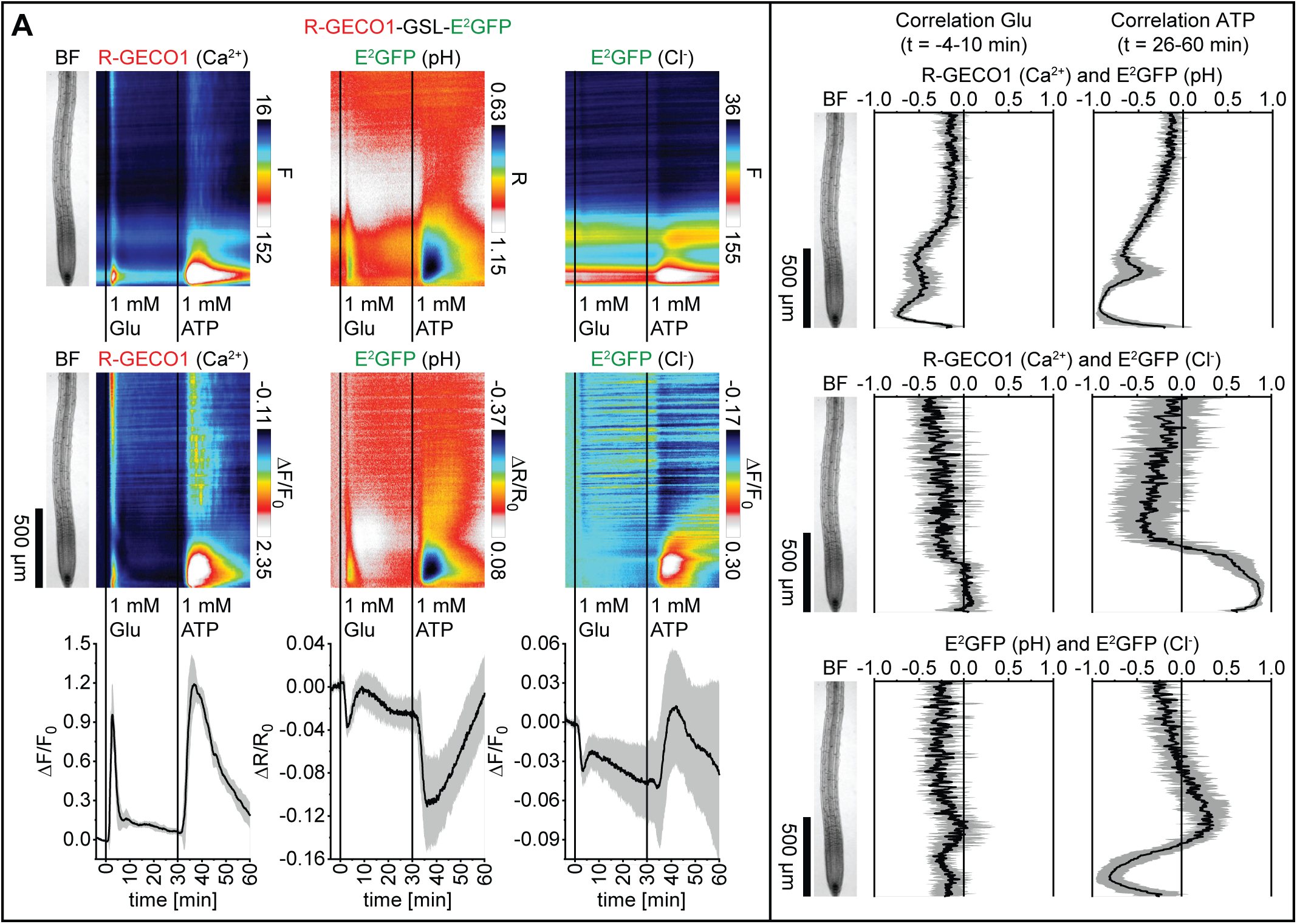
Application of glutamate triggers cytosolic Ca^2+^, H^+^ and Cl^−^ fluxes. Analyses of five-day-old roots of Arabidopsis expressing R-GECO1-GSL-E^2^GFP (Ca^2+^, pH and Cl^−^) in response to 1 mM glutamate (Glu; t = 0 min) and 1 mM ATP (t = 30 min; n = 7). Images were acquired for 64 min at a frame rate of 10 min^-1^. **(A)** Average vertical response profiles of (top) fluorescence emissions (F) or emission ratios (R), (middle) signal changes (ΔF/F_0_ or ΔR/R_0_) normalized to 4 min average baseline recordings, and (bottom) full image signal changes (mean ± SD). **(B)** Spatiotemporal Pearson correlation analyses (mean ± SD) of indicated GEFI responses during the application of glutamate (left; t = −4-10 min) or ATP (right; t = 26-60 min). An adjacent representative bright field (BF) root image is shown for orientation. A representative experiment is provided as Supplemental Movie 5. See also Supplemental Figures 3 and 4 for related experiments.

In order to increase the spatial resolution for pH measurements, R-GECO1-P2A-E^2^GFP was fused to the N-terminus of LOW TEMPERATURE INDUCED PROTEIN 6B (LTI6b) or VESICLE TRANSPORT V-SNARE 11 (VTI11). This enabled the targeting of E^2^GFP to the cytosolic side of the plasma membrane (LTI6b; Cutler et al., 2000) or the tonoplast (VTI11; Takemoto et al., 2018), while R-GECO1 remained in the cytosol and the nucleus (Supplemental Figures 3A and 4A). Compared to previous analyses (Figure 5), these indicators reported very similar Ca^2+^ and pH response patterns, irrespective of the subcellular localization of E^2^GFP. However, it appeared that Cl^−^ responses were more variable with noticeably higher Cl^−^ fluxes at the tonoplast (Supplemental Figures 3B and 4B). Note that R-GECO1-P2A-E^2^GFP-LTI6b expression induced more severe growth defects in Arabidopsis compared to the other GEFI lines (Supplemental Figure 5). Therefore, results obtained with this GEFI should be interpreted with caution. Taken together, R-GECO1-GSL-E^2^GFP enables the simultaneous monitoring of Ca^2+^, H^+^ and Cl^−^ fluxes, that in response to IAA, ATP and glutamate exhibited a remarkably high spatiotemporal overlap.

### Glutamate treatment induces cytosolic acidification without noticeable impact on the cytosolic redox state

To test whether glutamate has an impact on the cytosolic redox state, Arabidopsis seedlings expressing PA-17-P2A-Grx1-roGFP2 (pH and *E*_GSH_) or PA-17-P2A-roGFP2-Orp1 (pH and H_2_O_2_) were exposed to 1 mM glutamate and 100 µM H_2_O_2_ treatments as positive control. As observed before, glutamate triggered a biphasic cytosolic acidification, that prolonged during the 100 µM H_2_O_2_ response (Figures 6A left and 6B left, Supplemental Movies 6 and 7). Application of glutamate did not induce cytosolic redox changes. Whereas, 100 µM H_2_O_2_ treatments triggered a steep cytosolic oxidation that remained high for longer than 30 min (Figures 6A right and 6B right). During this response, the roGFPs indicated a cytosolic oxidation predominantly in epidermis and cortex cells of the elongation zone and above, with faster responses in upper root regions. Except of the epidermis, cells of the meristematic zone only slightly increased their redox state in response to H_2_O_2_ (Supplemental Movies 6 and 7). Altogether, these data indicate that 100 µM H_2_O_2_ treatments rapidly induce cytosolic oxidation, and that the root meristematic zone is less sensitive to this oxidative stress.

**Figure 6.**
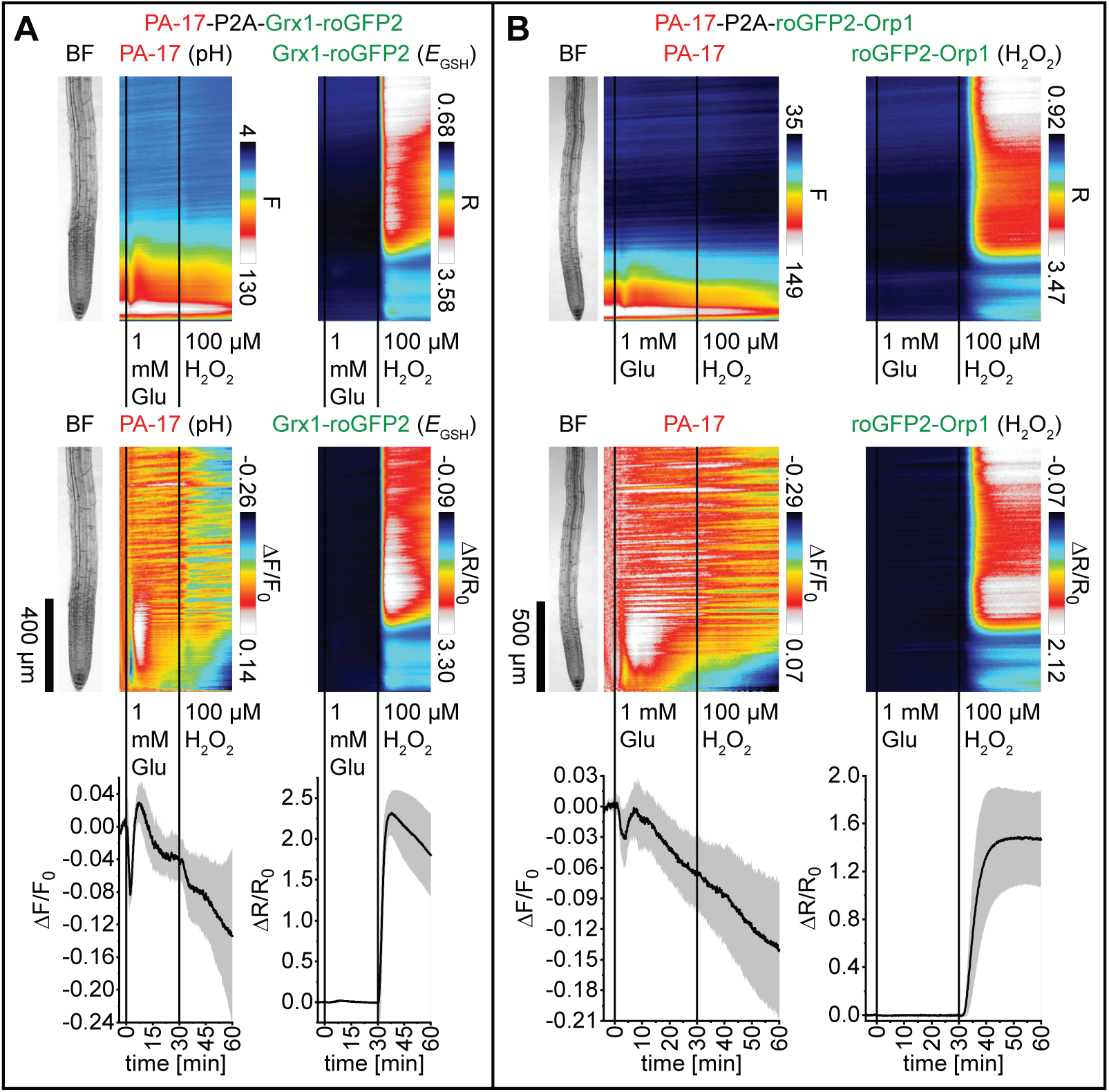
Application of Glutamate triggers a rapid cytosolic acidification without noticeable impact on the cytosolic redox state. Analyses of five-day-old roots of Arabidopsis expressing **(A)** PA-17-P2A-Grx1-roGFP2 (pH and *E*_GSH_; n = 6) and **(B)** PA-17-P2A-roGFP2-Orp1 (pH and H_2_O_2_; n = 8) in response to 1 mM glutamate (Glu; t = 0 min) and 100 µM H_2_O_2_ (t = 30 min). Images were acquired for 64 min at a frame rate of 10 min^-1^. Average vertical response profiles of (top) fluorescence emissions (F) or emission ratios (R) and (middle) signal changes (ΔF/F_0_ or ΔR/R_0_) normalized to 4 min average baseline recordings. An adjacent representative bright field (BF) root image is shown for orientation. (bottom) Full image signal changes (mean ± SD). Note that experiments in **(A)** and **(B)** were acquired at different magnifications. Representative experiments are provided as Supplemental Movies 6 and 7.

### H_2_O_2_ treatment triggers spatiotemporally overlapping but also distinct patterns of cytosolic oxidation and Ca^2+^ fluxes

Current models propose an interdependence of Ca^2+^- and ROS signaling (Gilroy et al., 2014; Steinhorst and Kudla, 2014). To investigate the spatiotemporal relationships of cytosolic Ca^2+^ and ROS signals, we first treated Arabidopsis seedlings expressing R-GECO1-P2A-Grx1-roGFP2 (Ca^2+^ and *E*_GSH_) or R-GECO1-P2A-roGFP2-Orp1 (Ca^2+^ and H_2_O_2_) with 20 and 100 µM H_2_O_2_. The *E*_GSH_ and H_2_O_2_ indicators responded to both treatments with similar patterns, albeit with increased signal changes in response to 100 µM H_2_O_2_ (Figures 7A right and 7B right, Supplemental Movies 8 and 9). Although 20 µM H_2_O_2_ treatments induced a detectable cytosolic oxidation, discernible Ca^2+^ signals were not observed (Figure 7, Supplemental Movies 8 and 9). In response to 100 µM H_2_O_2_ treatments, cytosolic oxidation preceded detectable Ca^2+^ signals. Although, both signals appeared to arrive from upper root regions, Ca^2+^ signals in the elongation zone exhibited a maximum response in the vasculature, whereas cytosolic oxidation was more pronounced in epidermis and cortex cells. Both signals exhibited a minimum response in the meristematic zone (Figure 7, Supplemental Movies 8 and 9). We conclude that these 2-In-1-GEFIs exhibit sufficient sensitivity for resolving the interrelation of cytosolic Ca^2+^ and ROS signals, which, in response to 100 µM H_2_O_2_ treatments, exhibit overlapping but not similar spatiotemporal response patterns. In addition, the roGFPs facilitate the detection of cytosolic oxidation in response to H_2_O_2_ below the threshold of Ca^2+^ channel activation.

**Figure 7.**
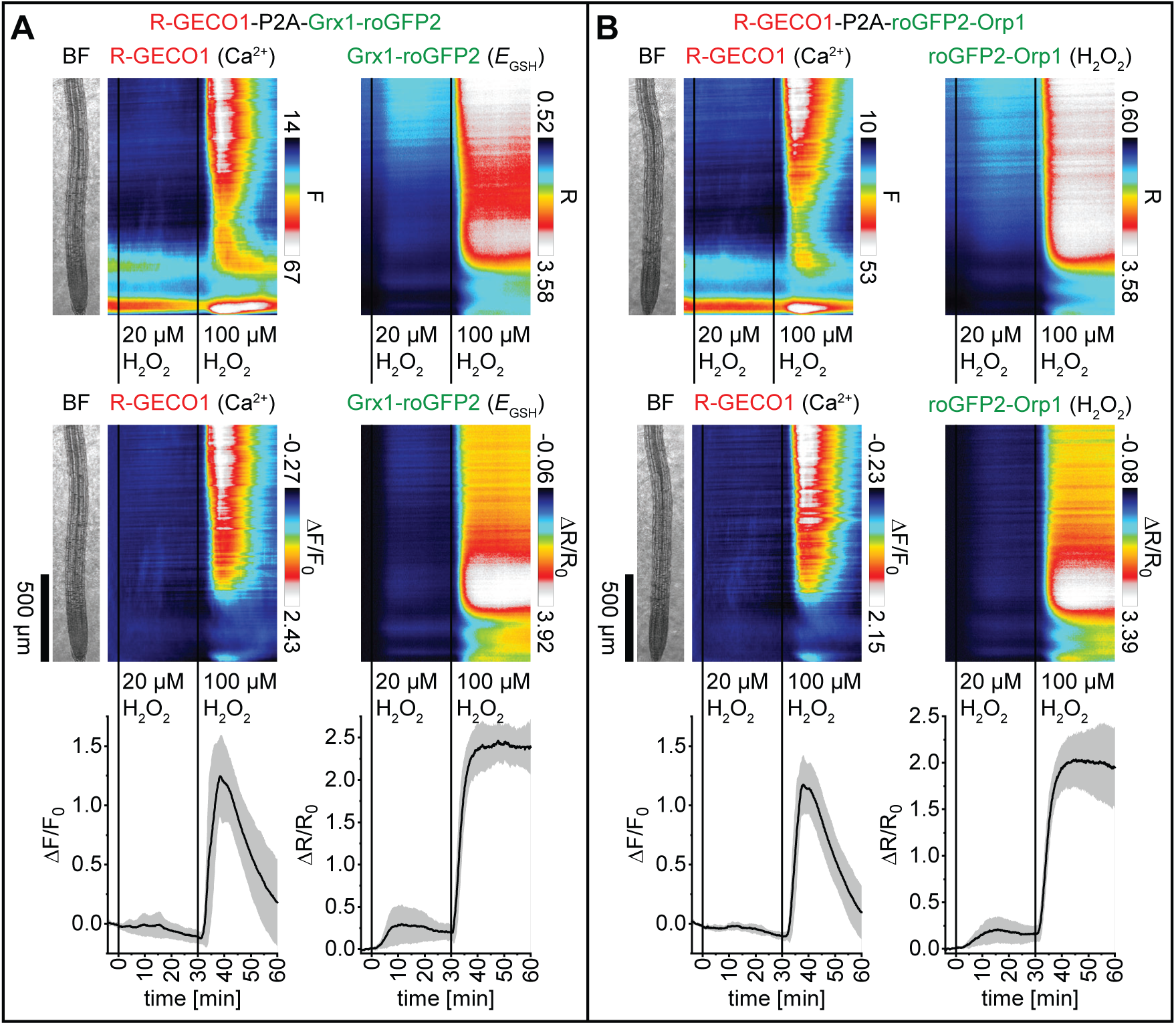
Application of H_2_O_2_ triggers overlapping but also distinct patterns of cytosolic oxidation and Ca^2+^ fluxes. Analyses of five-day-old roots of Arabidopsis expressing **(A)** R-GECO1-P2A-Grx1-roGFP2 (Ca^2+^ and *E*_GSH_; n = 8) and **(B)** R-GECO1-P2A-roGFP2-Orp1 (Ca^2+^ and H_2_O_2_; n = 8) in response to 20 µM H_2_O_2_ (t = 0 min) and 100 µM H_2_O_2_ (t= 30 min). Images were acquired for 64 min at a frame rate of 10 min^-1^. Average vertical response profiles of (top) fluorescence emissions (F) or emission ratios (R) and (middle) signal changes (ΔF/F_0_ or ΔR/R_0_) normalized to 4 min average baseline recordings. An adjacent representative bright field (BF) root image is shown for orientation. (bottom) Full image signal changes (mean ± SD). Representative experiments are provided as Supplemental Movies 8 and 9.

### ATP and AtPEP1 treatments trigger Ca^2+^, H^+^ and Cl^−^ fluxes, and a weak cytosolic oxidation

Extracellular ATP and the signaling peptide AtPEP1 function as damage-associated elicitors that trigger Ca^2+^ signals and ROS production (Song et al., 2006; Demidchik et al., 2009; Ma et al., 2014). To investigate the spatiotemporal relationships of these processes, Arabidopsis seedlings expressing R-GECO1 and roGFP2-Orp1 or Grx1-roGFP2 from individual expression cassettes located on one T-DNA, were subjected to 1 mM ATP and consecutive 100 µM H_2_O_2_ treatments as positive control. ATP triggered typical Ca^2+^ responses, but its effect on the cytosolic redox state was rather weak (Figure 8A, Supplemental Figure 6A). 100 µM H_2_O_2_ treatments induced Ca^2+^ fluxes and cytosolic oxidation as observed before (Supplemental Movies 10 and 12). Experiments using R-GECO1-P2A-roGFP2-Orp1 and R-GECO1-P2A-Grx1-roGFP2 revealed that 1 µM (At)PEP1 treatments induced Ca^2+^ signals that initiated in epidermis cells, followed by an overall Ca^2+^ burst, after which Ca^2+^ oscillations in the meristem appeared that proceeded to the vasculature and further shootward. However, PEP1 treatments had only weak effects on the cytosolic redox state (Figure 8B, Supplemental Figure 6B, Supplemental Movies 11 and 13). To better resolve the roGFP responses, the initial 30 min signal change response profiles were extracted from original data sets and calibrated to the same color scale. The data indicate a detectable cytosolic oxidation in response to glutamate, ATP and PEP1 that was however low compared to the 20 µM H_2_O_2_ response (Supplemental Figure 7). We also investigated the effect of PEP1 using R-GECO1-GSL-E^2^GFP. 1 µM PEP1 triggered a transient Ca^2+^, H^+^ and Cl^−^ influx that, during the initial 20 min of the response, exhibited a spatiotemporal overlap/correlation mainly in the meristematic- and elongation zone (Figures 9A and 9B, Supplemental Movie 14). Altogether, these experiments established that PEP1 triggers spatiotemporally overlapping Ca^2+^, H^+^ and Cl^−^ fluxes in roots. Whereas, the effect of PEP1 and ATP on the cytosolic redox state was below the threshold of ROS-induced Ca^2+^ signaling (Supplemental Figure 7).

**Figure 8.**
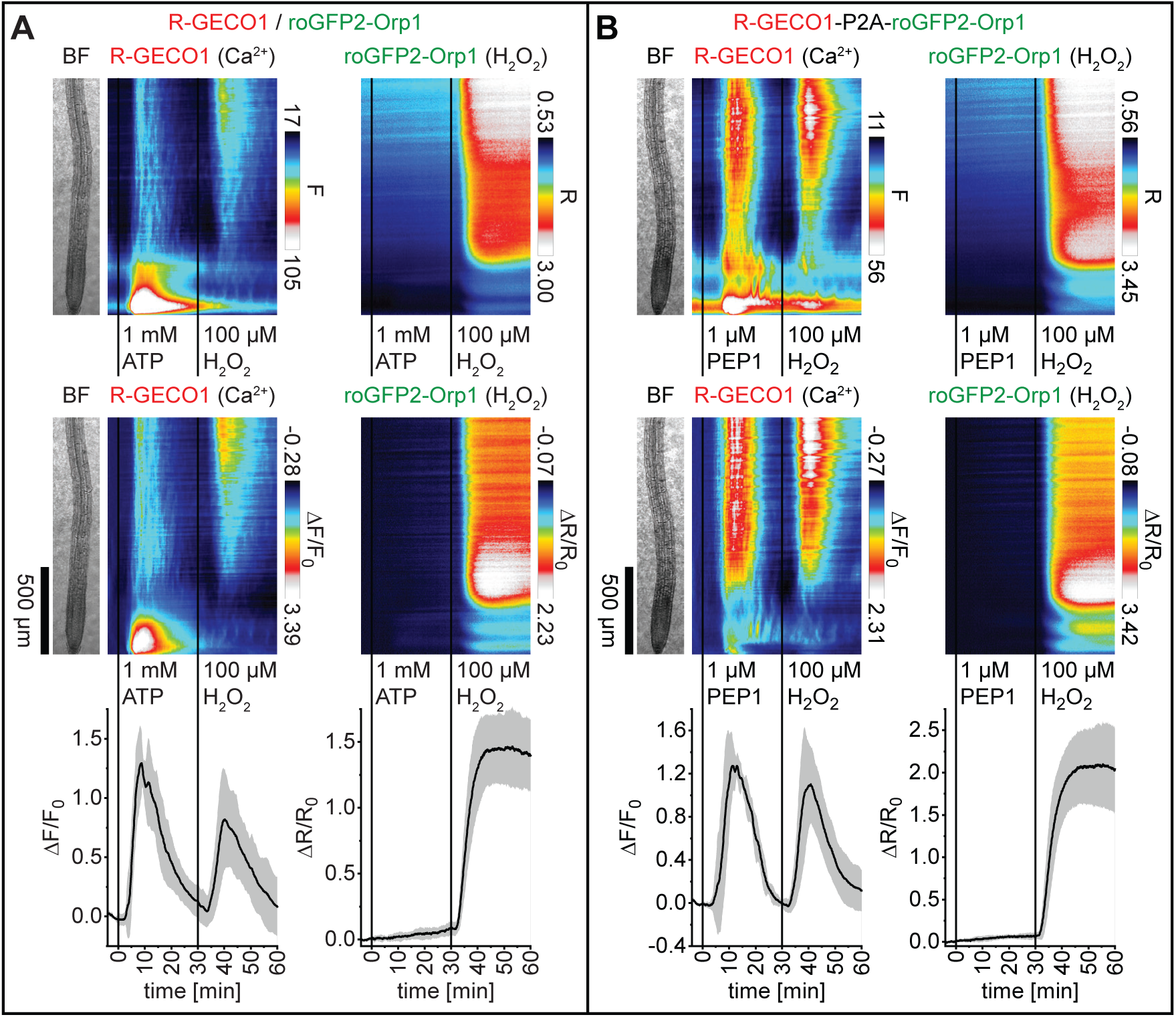
Cytosolic oxidation is only weakly affected by ATP and PEP1. Analyses of five-day-old roots of Arabidopsis expressing **(A)** R-GECO1 and roGFP2-Orp1 (Ca^2+^ and H_2_O_2_; n = 7) in response to 1 mM ATP and 100 µM H_2_O_2_, and **(B)** R-GECO1-P2A-roGFP2-Orp1 (Ca^2+^ and H_2_O_2_; n = 7) in response to 1 µM PEP1 and 100 µM H_2_O_2_. Average vertical response profiles of (top) fluorescence emissions (F) or emission ratios (R) and (middle) signal changes (ΔF/F_0_ or ΔR/R_0_) normalized to 4 min average baseline recordings. An adjacent representative bright field (BF) root image is shown for orientation. (bottom) Full image signal changes (mean ± SD). Representative experiments are provided as Supplemental Movies 10 and 11. See also Supplemental Figures 6 and 7 for related experiments.

**Figure 9.**
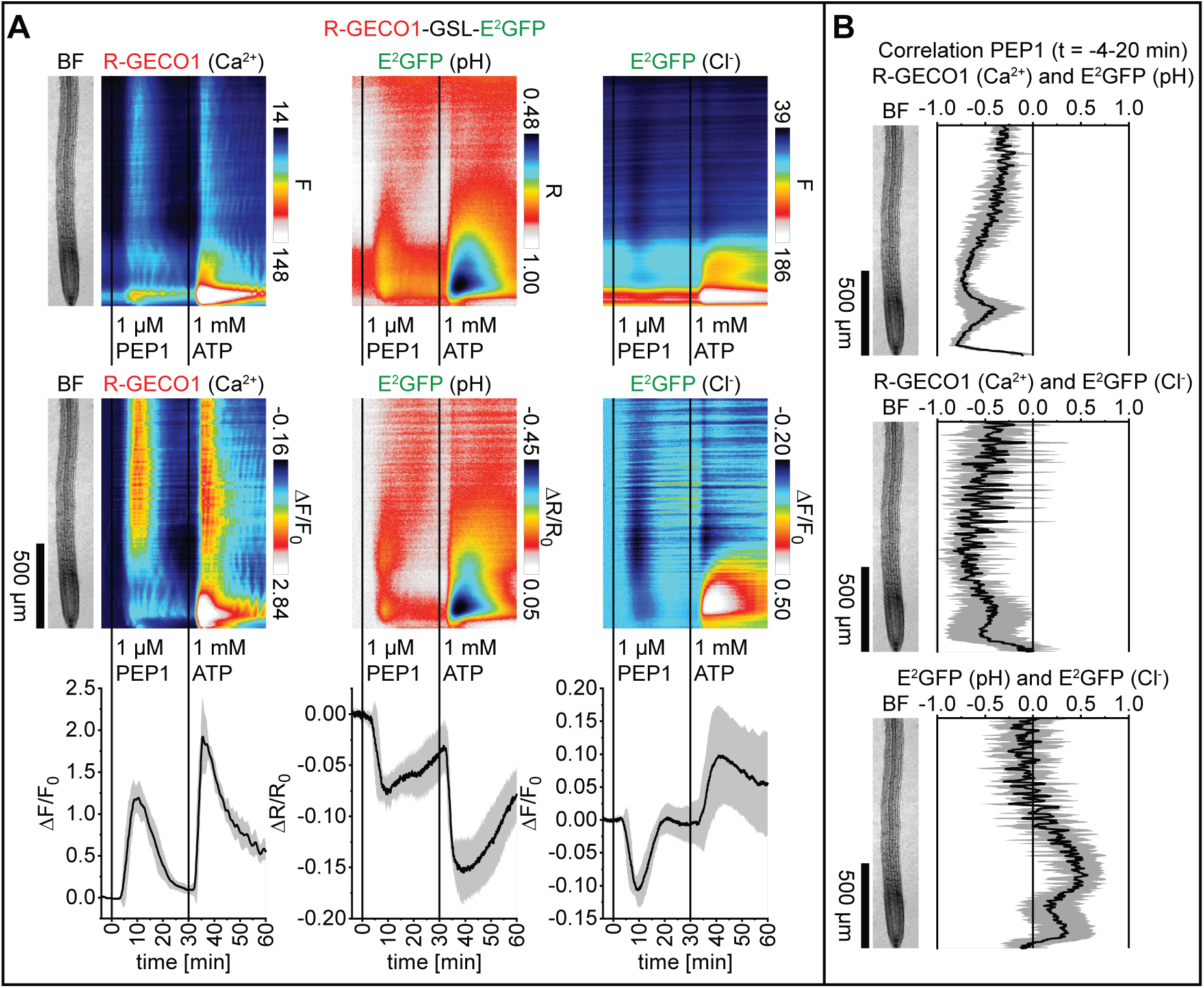
Application of PEP1 triggers Ca^2+^, H^+^ and Cl^−^ fluxes with high spatiotemporal overlap. **(A)** Analyses of five-day-old roots of Arabidopsis expressing R-GECO1-GSL-E^2^GFP (Ca^2+^, pH and Cl^−^; n = 6) in response to 1 µM PEP1 (t = 0 min) and 1 mM ATP (t= 30 min). Images were acquired for 64 min at a frame rate of 10 min^-1^. Average vertical response profiles of (top) fluorescence emissions (F) or emission ratios (R) and (middle) signal changes (ΔF/F_0_ or ΔR/R_0_) normalized to 4 min average baseline recordings. An adjacent representative bright field (BF) root image is shown for orientation. (bottom) Full image signal changes (mean ± SD). **(B)** Spatiotemporal Pearson correlation analyses (mean ± SD) of indicated GEFI responses during the application of PEP1 (t = - 4-20 min). A representative experiment is provided as Supplemental Movie 14.

### Glutathione disulfide (GSSG) treatments trigger rapid Ca^2+^, H^+^ and Cl^−^ fluxes that precede a slow-progressing cytosolic oxidation

GSSG is known to trigger cytosolic Ca^2+^ signals (Gomez et al., 2004) and to directly oxidize Grx1-roGFP2 (Gutscher et al., 2008). We sought to resolve the spatiotemporal relations of these responses in Arabidopsis seedlings expressing R-GECO1-P2A-Grx1-roGFP2. Although 1 mM GSSG-induced Ca^2+^ signals were variable, they appeared to arrive from upper root regions and travelled towards the root tip, followed by a second Ca^2+^ burst in the vasculature and oscillations in the meristematic- and elongation zone (Figure 10A left, Supplemental Movie 15). After the initial Ca^2+^ signal reached the root tip, in this region a cytosolic oxidation was observed that slowly progressed towards the elongation zone, where oscillation became visible (Figure 10A right, Supplemental Movie 15). Note that the Ca^2+^ and *E*_GSH_ oscillations were shifted in phase (Supplemental Movie 15). Additional experiments using R-GECO1-GSL-E^2^GFP revealed that GSSG treatments also induced H^+^ and Cl^−^ influx, exhibiting the most noticeable spatiotemporal overlap with Ca^2+^ signals in the meristematic- and early elongation zone during the initial 20 min of the GSSG response (Figures 10B and 10C, Supplemental Movie 16). In summary, GSSG treatments trigger Ca^2+^ signals, cytosolic acidification and Cl^−^ influx that precede a cytosolic oxidation.

**Figure 10.**
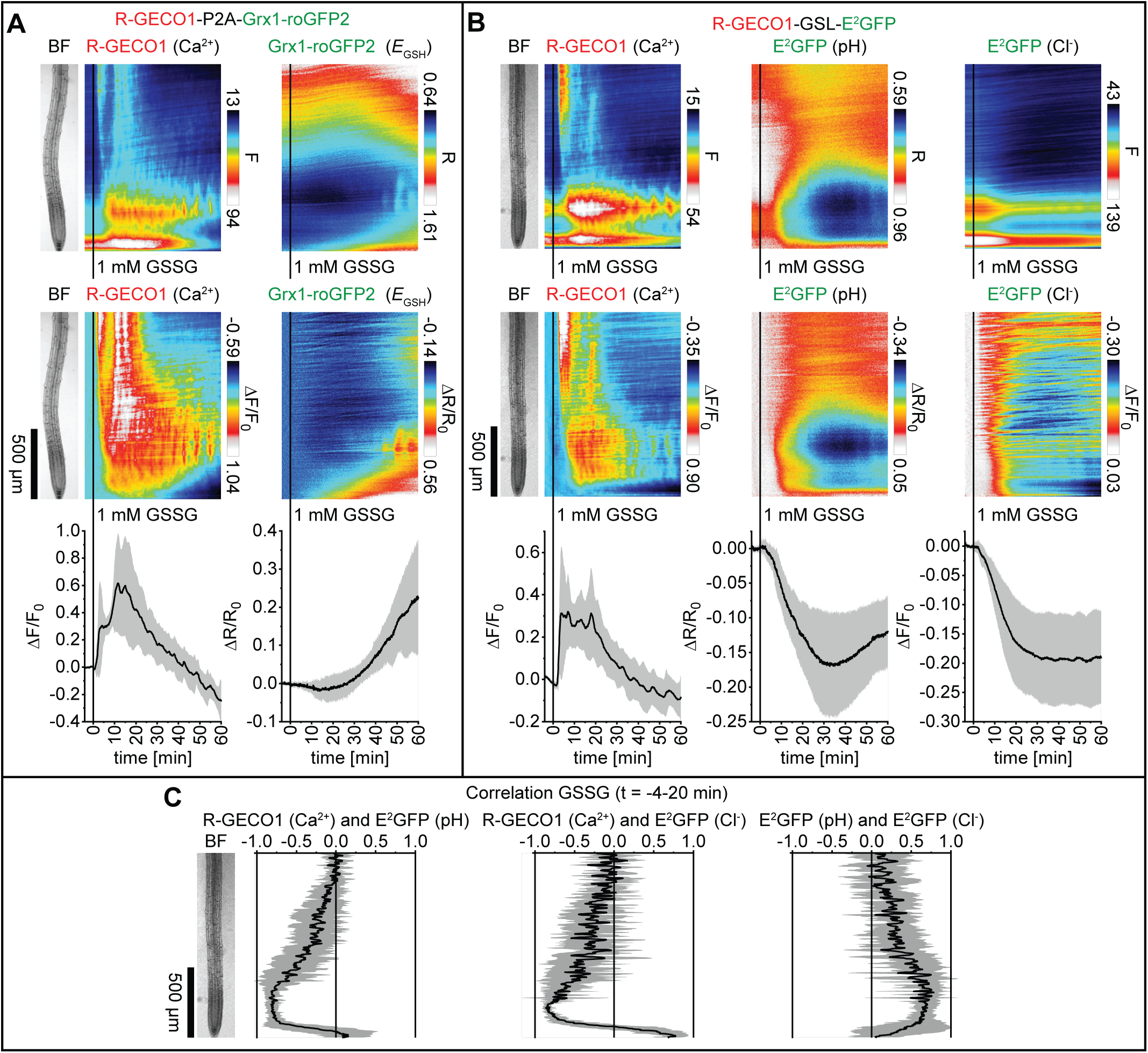
GSSG-triggered Ca^2+^-, H^+^- and Cl^−^ influx precedes cytosolic oxidation. Analyses of five-day-old roots of Arabidopsis expressing **(A)** R-GECO1-P2A-Grx1-roGFP2 (Ca^2+^ and *E*_GSH_; n = 5) and **(B)** R-GECO1-GSL-E^2^GFP (Ca^2+^, pH and Cl^−^; n = 6) in response to 1 mM GSSG (t = 0 min). Images were acquired for 64 min at a frame rate of 10 min^-1^. Average vertical response profiles of (top) fluorescence emissions (F) or emission ratios (R) and (middle) signal changes (ΔF/F_0_ or ΔR/R_0_) normalized to 4 min average baseline recordings. An adjacent representative bright field (BF) root image is shown for orientation. (bottom) Full image signal changes (mean ± SD). **(C)** Spatiotemporal Pearson correlation analyses (mean ± SD) of indicated GEFI responses during the application of GSSG (t = −4-20 min; data from **(B)**). Representative experiments are provided as Supplemental Movies 15 and 16.

## DISCUSSION

### Optimization of ABA indicators in HEK293T cells

Optimization procedures of FRET-based indicators usually comprise the testing of FRET-pair-, sensory domain- and linker variants (Okumoto et al., 2012; Hochreiter et al., 2015). Although such testing has been performed on ABACUS, the linkers between the sensory domain and attached fluorescent proteins remained invariant (Jones et al., 2014). Early optimizations of ABAleon focused on sensory domain modifications that led to the development of ABAleon2.15 with improved stereospecificity for (+)-ABA (Waadt et al., 2014). Using the HEK293T cell system, we have developed ABAleonSD1-3L21 that exhibits an improved signal-to-noise ratio compared to ABAleon2.15 (Figure 1B). HEK293T cells are a convenient system for GEFI screenings, because they can be easily transfected, cultivated and analyzed in a plate reader (Tian et al., 2009). HEK293T cells contain neglectable ABA levels, and are therefore well suited for ABA indicator screenings with a potential to facilitate the heterologous characterization of ABA transporters using ABA indicators. Successful characterizations of plasma membrane proteins in HEK293T cells has been demonstrated for RBOHs and CYCLIC NUCLEOTIDE-GATED ION CHANNEL (CNGC)-type Ca^2+^ channels (Gao et al., 2016; Han et al., 2019). The differences of ABAleon characteristics between HEK293T cell and in vitro analyses might be due to a lower stability of ABAleon2.15 in HEK293T cells. Similar issues have been reported for Ca^2+^ indicators (Tian et al., 2009). However, in vitro characterizations of ABA indicators are time-consuming, and screenings using *E. coli* are not practical due to a likely even lower protein stability in this system (Jones et al., 2014; Waadt et al., 2014). Note that previously measured properties of ABAleon2.15 (ΔR_(max)_/R_0_ ∼ −0.10 and k’_d_ ∼ 500 nM; Waadt et al., 2014) were different compared to results in Figures 1C and 1F. Here we used a faster sandwich-tag purification procedure with subsequent characterization of freshly purified proteins that might give more reliable results.

ABAleon2.15 and ABAleonSD1-3L21 exhibited similar ABA responses in Arabidopsis. ABACUS1-2µ responded slower to ABA and did not resolve the ABA gradient in roots (Figure 2, Supplemental Figure 2), probably due to the lower ABA affinity (Jones et al., 2014). However, this indicator might have advantages for the analyses of ABA dynamics in the root tip, the root-hypocotyl junction and in guard cells, where ABAleons are close to saturation (Figure 2; Waadt et al., 2015). In the future, optimization of ABA indicators will require the development of alternative sensory domains and the investigation of alternative biosensor designs.

### 2-In-1-GEFIs facilitate simultaneous multiparametric analyses

Multiplexed live imaging in plants has been performed via the combination of GEFIs with fluorescent dyes, the use of GEFIs in parallel experiments, or through dual-expression of Ca^2+^ indicators (Monshausen et al., 2007, 2009, 2011; Loro et al., 2012; Schwarzländer et al., 2012; Ngo et al., 2014; Keinath et al., 2015; Behera et al., 2018; Kelner et al., 2018; Wagner et al., 2019). However, GEFI-based simultaneous analyses of two signaling compounds has been established in Arabidopsis only for Ca^2+^ and ABA (Waadt et al., 2017). Because most GEFIs are FRET-or green fluorescent protein-based, simultaneous multiparametric analyses have become possible through the development of red fluorescent protein-based indicators for Ca^2+^, redox/H_2_O_2_ and pH (Bilan and Belousov, 2017; Martynov et al., 2018; Walia et al., 2018). Yet, except for R-GECO1, their application in plants is rare. Here, we introduced the intensiometric red fluorescing pH indicator (P)A-17 (Shen et al., 2014), which is well suited for pH analyses in Arabidopsis with similar responsiveness compared to the ratiometric E^2^GFP (Figures 3B left, 4A middle and 5A middle).

As the generation of stable transgenic organisms is time-consuming, simultaneous expression of GEFIs, or the generation of dual-sensing GEFIs, is advantageous. Moreover, the latter approach minimizes epigenetic silencing effects, often observed in lines carrying multiple transgenes. Dual-sensing GEFIs have been developed for pH and Cl^−^ (ClopHensor; Arosio et al., 2010) and for phosphatidylinositol 3,4,5-trisphosphate localization and H_2_O_2_ concentration (PIP-SHOW; Mishina et al., 2012). For the generation of our 2-In-1-GEFIs we were inspired by ClopHensor and the incorporated E^2^GFP that we fused with R-GECO1 in analogy to R-GECO1-GSL-mTurquoise (Waadt et al., 2017). Note that recent studies indicated that ClopHensor/E^2^GFP might also respond to NO_3_^−^ (https://doi.org/10.1101/716050). Because our imaging buffer contained 5 mM Cl^−^ and the microscope-dish agarose media contained 4 mM NO_3_^−^, the observed E^2^GFP responses likely depended on both anion species. In contrast to R-GECO1-GSL-E^2^GFP, the other 2-In-1-GEFIs have been linked via the self-cleaving P2A-peptide, which enables efficient cleavage in Arabidopsis (Burén et al., 2012; Supplemental Figures 3 and 4). In addition, P2A-based 2-In-1-GEFIs exhibit similar responses compared to indicators expressed from separate expression cassettes (Figure 8, Supplemental Figure 6). Because only one expression cassette is used, P2A-linked GEFIs can be more easily screened at the microscope and are less prone to unwanted silencing effects. Our work established several 2-In-1-GEFIs based on the P2A-linkage, which is applicable to any eukaryotic system (Kim et al., 2011).

### Ca^2+^, H^+^ and anion fluxes exhibit a high spatiotemporal overlap

Previous work established that mechanical stimulation, wounding, ATP and auxin simultaneously induce Ca^2+^ and H^+^ fluxes (Monshausen et al., 2009, 2011; Behera et al., 2018). We found that, in addition to auxin and ATP, also glutamate, PEP1 and GSSG trigger Ca^2+^, H^+^ and anion fluxes with high spatiotemporal overlap (Figures 4, 5, 9 and 10). The linkage of Ca^2+^ and H^+^ fluxes may depend on H^+^ pumps and Ca^2+^/H^+^-coupled transport via CATION/PROTON EXCHANGERS (CAXs) or Ca^2+^-ATPases (Bonza and De Michelis, 2011; Pittman and Hirschi, 2016). However, knowledge about their role in Ca^2+^ signaling is fragmentary, probably due to functional overlap of gene family members (Behera et al., 2018). Simultaneous Ca^2+^ and H^+^ fluxes in response to auxin are mediated by the auxin/H^+^-symporter AUXIN RESISTANT 1 (AUX1) and the Ca^2+^ channel CNGC14 that are functionally coupled (Shih et al., 2015; Dindas et al., 2018). Since the activation of plasma membrane H^+^-ATPases is coupled to AUX1 (Inoue et al., 2016), this could explain the subsequent H^+^-efflux.

Mechanical stimulation-induced Ca^2+^ and H^+^ fluxes depend on the RECEPTOR-LIKE KINASE (RLK) FERONIA, which acts as a receptor for RAPID ALKALIZATION FACTOR (RALF) peptides (Haruta et al., 2014; Shih et al., 2014; Stegmann et al., 2017). Several RLKs, including the ATP receptor DOES NOT RESPOND TO NUCLEOTIDES 1 (DORN1) and PEP RECEPTORS (PEPRs), can induce Ca^2+^ signals, apoplastic alkalization (coupled to cytosolic acidification), and ROS production (Qi et al., 2010; Choi et al., 2014; Ma et al., 2014; Seybold et al., 2014; Haruta et al., 2015; Chen et al., 2017; Kimura et al., 2017). The effect of DORN1 and PEPRs on anion efflux was observed in guard cells during stomatal closure (Chen et al., 2017; Zheng et al., 2018). Our analyses revealed that PEP1 induced a transient anion influx along the entire imaged root axis. Extracellular ATP triggered anion influx in the differentiation- and elongation zone, but efflux in the meristematic zone (Figure 9). Early research revealed that cytosolic but not extracellular ATP is required for adenine nucleotide activation of R-type anion channels (Hedrich et al., 1990) and protein kinase-mediated activation of S-type anion channels (Schmidt et al., 1995). It is likely that ATP-triggered Ca^2+^ signals activate Ca^2+^-dependent protein kinases required for the activation of anion channels (Mori et al., 2006). The differences in PEP1- and ATP-induced anion fluxes in roots might be due to the distinct Ca^2+^ signatures observed in the meristematic zone (Figure 9). In the future, it will be interesting to discriminate the differences in anion-flux regulation between roots and guard cells.

### On the interdependence of Ca^2+^ and ROS signaling

The interdependence of Ca^2+^ and ROS signaling has been extensively discussed (Gilroy et al., 2014; Steinhorst and Kudla, 2014). In the context of long-distance and systemic signaling, current models propose that Ca^2+^ signals trigger the ion channel TPC1 for signal amplification. Ca^2+^ signal propagation occurs via plasmodesmata or Ca^2+^-dependent activation of RBOHs. RBOH-derived extracellular ROS propagate to adjacent cells to activate plasma membrane localized Ca^2+^ channels (Evans et al., 2016; Choi et al., 2016). Ca^2+^-dependent activation of RBOHs is well established (Kadota et al., 2015; Han et al., 2019). However, whether RBOH-dependent ROS contribute to Ca^2+^ channel activation, has only been inferred from pharmacological- and genetic analyses (Kwak et al., 2003; Evans et al., 2016). In Arabidopsis guard cells, hyperpolarization-activated Ca^2+^-permeable channels can be activated by 50 µM H_2_O_2_ (Pei et al., 2000). In *Vicia faba* guard cells such channels exhibit a half response at 76 µM H_2_O_2_ (Köhler et al., 2003). Analyses in root epidermis cells revealed a Ca^2+^ channel activation by 10 µM H_2_O_2_ in the elongation zone and by 1 mM H_2_O_2_ in the maturation zone (Demidchik et al., 2007). The threshold concentrations of ROS required to activate Ca^2+^ channels may depend on the cell type, the location (apoplast or cytosol) and the chemical nature of ROS (Demidchik et al., 2007).

Our analyses revealed that in Arabidopsis roots 20 µM extracellular H_2_O_2_ triggered a detectable cytosolic oxidation, but no considerable Ca^2+^ signals (Figure 7). On the other hand, glutamate, ATP and PEP1, which efficiently trigger Ca^2+^ signals, induced a cytosolic oxidation rather below this threshold (Supplemental Figure 7). These data are consistent with a slow progressing cytosolic oxidation in response to the pathogen-associated molecular pattern flagellin fragment flg22 (Nietzel et al., 2019). Whether such cytosolic oxidation is Ca^2+^ dependent, requires further experimentation. However, compared to 20 µM H_2_O_2_ responses, our data suggest that glutamate-, ATP- and PEP1-induced cytosolic oxidation is not sufficient to trigger root Ca^2+^ channels (Supplemental Figure 7). We hypothesize that the observed ROS dependence of Ca^2+^ signaling may be indirectly linked to the impact of ROS on the cell wall, which binds considerable amounts of Ca^2+^ in Ca^2+^-pectate cross-linked complexes (Hepler et al., 2010; Peaucelle et al., 2012; Kärkönen and Kuchitsu, 2015). Such a model would be consistent with a rather slow H_2_O_2_ activation of Ca^2+^ channels in patch clamp analyses (20-60 min; Demidchik et al., 2007). Another possibility would be that a signaling component triggers both, Ca^2+^ and ROS signaling. The BOTRYTIS-INDUCED KINASE 1 (BIK1) could be a good candidate for such a mechanism (Kadota et al., 2014; Li et al., 2014; Kimura et al., 2017; Tian et al., 2019).

### Conclusions

Our work established 2-In-1-GEFI-based simultaneous multiparametric in vivo analyses of signaling compounds in Arabidopsis. Using the 2-In-1-GEFIs, we observed that in roots ABA does not trigger rapid Ca^2+^ or pH changes. Whereas, auxin, glutamate, ATP, PEP1 and GSSG induce Ca^2+^, H^+^ and anion fluxes with high spatiotemporal overlap (Figure 11, Supplemental Figure 8). These results suggest an interdependence and coordination of ion fluxes that need to be dissected in future research. Findings that glutamate-, ATP- and PEP1-induced cytosolic oxidation is below the threshold required for triggering Ca^2+^ channels argue against the current model of a ROS-assisted Ca^2+^ signal propagation mechanism (Evans et al., 2016). We hypothesize that ROS may have an indirect effect on Ca^2+^ signaling. Overall, 2-In-1-GEFI-based imaging will allow for high resolution in vivo analyses of signaling processes beyond the model plant Arabidopsis.

**Figure 11.**
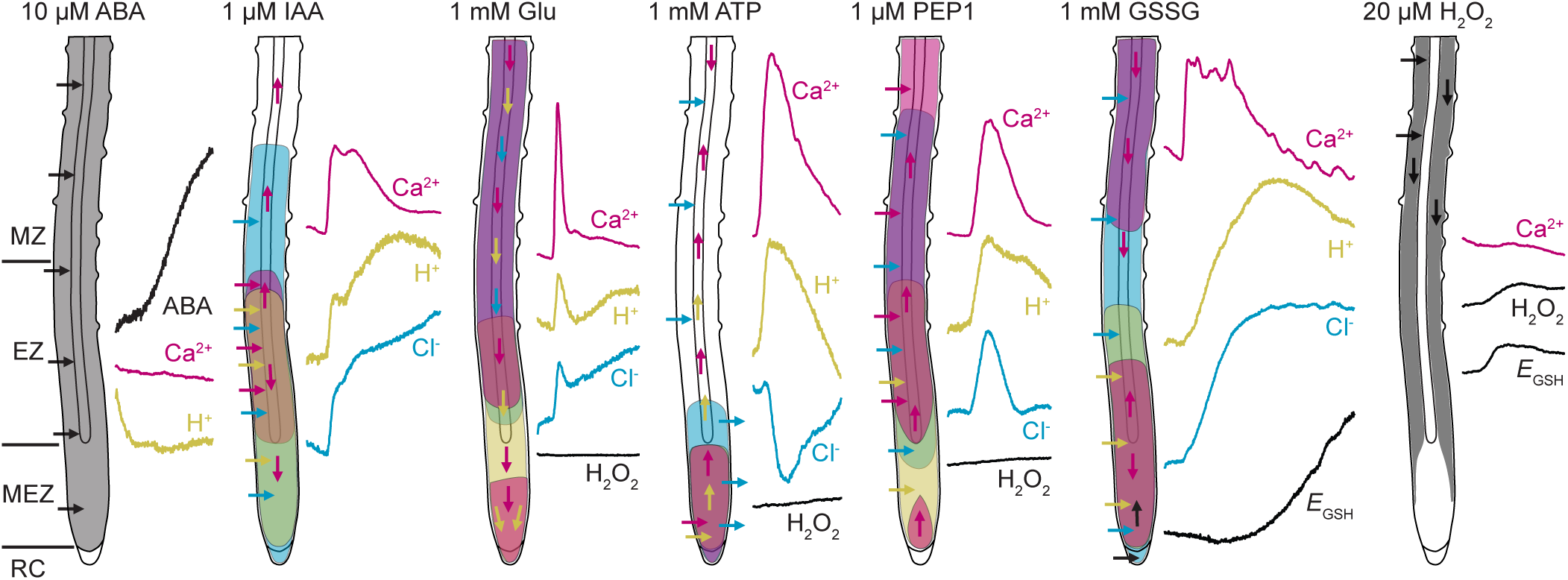
Schematic model of observed ABA, Ca^2+^, H^+^, Cl^−^ and redox changes in roots. ABA treatment and uptake did not induce rapid fluxes of Ca^2+^ or H^+^. Whereas, IAA, glutamate (Glu), ATP, PEP1 and GSSG triggered Ca^2+^, H^+^ and Cl^−^ fluxes with high spatiotemporal overlap. For comparison of the obtained data see also Supplemental Figure 8. Compared to 20 µM H_2_O_2_ and 1 mM GSSG, redox changes in response to glutamate, ATP and PEP1 were very low and below the threshold required to trigger ROS-induced Ca^2+^ signaling. Regions with highest response are color-coded according to the adjacent flux curves that were taken from the corresponding main figures (ABA, H_2_O_2_ and *E*_GSH_, black, Ca^2+^, magenta; H^+^, yellow; Cl^−^, cyan). For a better illustration of H^+^ and Cl^−^ influx the PA-17 and E^2^GFP response curves were inverted. Arrows indicate the direction of the corresponding fluxes. RC, root cap; MEZ, meristematic zone; EZ, elongation zone; MZ, maturation zone.

## METHODS

### Generation of plasmids

Oligonucleotides used for cloning procedures (Supplemental Data Set 1A) were obtained from Eurofinsgenomics. Plasmids (Supplemental Data Set 1B) were constructed using classical cloning procedures and the GreenGate system (Lampropoulos et al., 2013) utilizing enzymes from Thermo Fisher Scientific. Arabidopsis codon-optimized DNA fragments of PmTurquoise and PA-17 were designed using GeneArt^TM^ gene synthesis (Thermo Fisher Scientific). Expression of GEFIs in *Arabidopsis thaliana* Col-0 was carried out utilizing the promoter of a ubiquitous and highly expressed reference gene *ASPARTIC PROTEASE A1* (*APA1*, AT1G11910) that was chosen based on searches using Genevestigator (Hruz et al., 2008). The expression cassette also included the terminator of the *HEAT SHOCK PROTEIN 18.2* (*HSP18.2*) gene (AT5G59720; Nagaya et al., 2010; Waadt et al., 2014).

### Optimization of ABA indicators in HEK293T cells

Transformation and cultivation of HEK293T cells was performed as described previously (Ogasawara et al., 2008; Zhang et al., 2018). Spectral characteristics of ABA indicators were recorded in Greiner flat bottom 96-well plates (Greiner BIO-ONE) using a TECAN Safire plate reader (TECAN) operated by the XFLUOR4.51 software with the following parameters: fluorescence emission scan bottom mode; excitation wavelength 440 nm, bandwidth 12.5 nm; emission wavelength scan from 460-600 nm, bandwidth 10 nm; gain 100-115; flashes 10; integration time 40-60 µs; temperature 37 °C. HEK293T cells were kept in Hanks Balanced Salt Solution (HBSS; Thermo Fisher Scientific) and fluorescence emission spectra were recorded before (t_0_) and 60 min (t_60_) after exchange of solution to either HBSS with 100 µM (±)-ABA (Merck) and 0.1 % EtOH (treatment) or HBSS with 0.1 % EtOH (solvent control). ABA indicator emission ratios (R) were calculated as average emission at 518-538 nm divided by average emission at 470-490 nm after subtraction of the non-transfected HEK293T cell background emission spectrum. Emission ratio change (ΔR/R_0_) was calculated as [R(t_60_)-R(t_0_)]/R(t_0_). Experiments were performed in triplicates.

### Purification and in vitro characterization of ABAleons

BL21-CodonPlus (DE3)-RIL cells transformed with pET28-6xHis-ABAleon-(P)StrepII constructs were shaken at 150 rpm and 37 °C in 2x 1 L Luria Broth (LB) media supplemented with 50 µg mL^-1^ kanamycin and 30 µg mL^-1^ chloramphenicol. At an optical density (OD_600_) of 0.5, 1 mM Isopropyl Δ-D-1-thiogalactopyranoside (IPTG; Carl Roth) was added and protein expression was conducted in a shaking incubator at 24 °C for 6 h. Cultured cells were harvested by several centrifugation steps at 4 °C and 4000 rpm and bacterial pellet was flash frozen in liquid N_2_ and stored at - 80 °C.

The bacterial pellet was thawed on ice and resuspended in 30 mL lysis buffer (1x PBS [137 mM NaCl, 2.7 mM KCl, 10 mM Na_2_HPO_4_, 1.8 mM KH_2_PO_4_], 10 mM imidazole [Merck], 1x Roche protease inhibitor EDTA-free, 1 mM Phenylmethylsulfonyl fluoride [PMSF; Carl Roth] and 1 mg mL^-1^ lysozym [VWR], pH 7.4). After 40-60 min incubation on ice, cells were disrupted through microtip-based sonication (25 % amplitude, 21x 20 s) and cell debris were removed by centrifugation (2x 30 min, 20000 g, 4 °C) and filtering through 0.45 µm syringe filters (Merck).

6x-His purification was conducted in 20 mL gravity columns (VWR) loaded with 4 mL HisPur™ Ni-NTA resin (Thermo Fisher Scientific). After binding of proteins to the Ni-NTA resin, columns were washed 5x with 10 mL His-wash buffer (1x PBS, 25 mM imidazole, pH 7.4) and proteins were eluted in 3x 2 mL His-elution buffer (1x PBS, 250 mM imidazole, pH 7.4). Purified proteins were then loaded onto a 20 mL gravity column supplemented with 3 mL 50 % Strep Tactin Superflow (IBA). After 4x washing with 7.5 mL SII-wash buffer 1 (30 mM Tris/HCl pH 7.4, 250 mM NaCl) and 3x washing with 7.5 mL SII-wash buffer 2 (30 mM Tris/HCl pH 7.4, 250 mM NaCl, 10 mM MgCl_2_, 1 mM MnCl_2_), proteins were eluted in 3x 1.5 mL SII-elution buffer (30 mM Tris/HCl pH 7.4, 250 mM NaCl, 10 mM MgCl_2_, 1 mM MnCl_2_, 2.5 mM desthiobiotin [IBA]) and concentrated to a final volume of about 1 mL using Amicon Ultra-4 30 K filters (Merck). Purity of proteins was analyzed by SDS-PAGE using 10 % Mini-PROTEAN® TGX™ Precast Gels (BioRad) and InstantBlue staining (Expedeon). In a similar procedure, protein yield was calculated according to a bovine serum albumin standard curve.

For vitro calibration, a 100 mM (+)-ABA (TCI) stock solution dissolved in 100 % EtOH was used for an ABA dilution series in SII-wash buffer 2 and 0.2 % EtOH. 10 µL of each ABA dilution were added to 3 wells of black flat bottom µclear® 96-well plates (Greiner BIO-ONE) containing 90 µL of ∼ 1.1 µM ABAleon protein, diluted in SII-wash buffer 2, or to 90 µL SII-wash buffer 2 alone as background control. Fluorescence emission spectra were recorded after 20 min incubation at room temperature in the dark using a TECAN Infinite M1000 plate reader (TECAN) operated by the i-control 1.10.4.0 software with the following parameters: fluorescence emission scan bottom mode; excitation wavelength 440 nm, bandwidth 10 nm; emission wavelength scan from 460-650 nm, bandwidth 10 nm; gain 98-104, flashes 10 at 100 Hz, integration time 60 µs, temperature 21-22 °C. ABA-dependent ABAleon emission ratios (R) were calculated as described above. Maximum emission ratio change (ΔR_(max)_/R_0_) was calculated as [R(at 20 µM ABA)-R(at 0 µM ABA)]/R(at 0 µM ABA). Apparent ABA affinities (k’_d_; EC50) of ABAleons were calculated by fitting the emission ratio values of all three replicates to a 4-parameter logistic function using OriginPro 2018 (OriginLab Corporation).

### Generation of transgenic Arabidopsis plants expressing GEFIs

*Agrobacterium* strain ASE containing the pSOUP helper plasmid and the respective plant expression vectors (Supplemental Data Set 1B) were used for transformation of *Arabidopsis thaliana* ecotype Col-0 by floral dip (Clough & Bent, 1998) to generate the transgenic lines listed in Supplemental Data Set 1C. Seeds of transformed plants were surface sterilized for 10-15 min in 70 % EtOH, washed three times with 100 % EtOH and sowed on half-strength Murashige & Skoog (0.5 MS) media (Duchefa) supplemented with 5 mM MES-KOH pH 5.8, 0.8 % phytoagar (Duchefa) and 10 µg mL^-1^ Glufosinate-ammonium or 25 µg mL^-1^ hygromycin B (Merck) for herbicide selection. After 3-6 days of stratification in the dark at 4 °C, transgenic plants were grown for six-days in a growth room (16 h day/8 h night, 22 °C, 65 % relative humidity, photon fluence rate 100 µmol m^-2^ s^-1^). Positive transformants were then transferred to herbicide-free 0.5 MS media-containing petri dishes. After one day recovery, GEFI expression was confirmed by visual inspection at a Zeiss Discovery.V20 fluorescence stereo microscope equipped with GFP, YFP and RFP filters and a Plan S 1.0x FWD 81 mm lens. Approximately 40 herbicide resistant and fluorescing seedlings were then transferred to round 7 cm pots containing classic soil (Einheitserde) and grown until seed ripening in the growth room. ABAleon expressing plants were covered with a plastic lid and grown in a Conviron CMP6010 growth chamber (16 h day/8 h night, 20 °C, 65 % relative humidity, photon fluence rate 150 µmol m^-2^ s^-1^). To confirm proper GEFI expression, compare GEFI fluorescence emissions and avoid silencing effects in next generations, one leaf of each individual about three-week-old plant was examined at a confocal laser scanning microscope Leica SP5 II equipped with a HCX PL APO CS 20.0 x 0.7 IMM UV objective (Leica Microsystems) using emission and excitation settings listed in Supplemental Data Set 1D. For each construct, at least two transgenic lines with highest GEFI expression, proper 3:1 segregation in the 2^nd^ generation and least silencing were selected for further propagation. One line, indicated with (microscope) in Supplemental Data Set 1C, was used for microscopic experiments.

### Phenotypic characterization of GEFI lines

Seeds were surface sterilized and sown on 0.5 MS media supplemented with 5 mM MES-KOH pH 5.8 and 0.8 % phytoagar. Seven-day-old seedlings grown in the growth room (16 h day/8 h night, 22 °C, 65 % relative humidity, photon fluence rate 100 µmol m^-2^ s^-1^) were transferred to soil in single pots and further grown until 28-days-old. Pictures from 9-12 plants per genotype were acquired from the top and rosette area values were obtained using the Rosette Tracker Fiji plugin (De Vylder et al., 2012).

### Microscopic analyses

Seeds of GEFI expressing lines were surface sterilized and sown in four horizontal rows on square petri dishes containing LAK media (Barragán et al., 2012; 1 mM KH_2_PO_4_, 2 mM Ca(NO_3_)_2_, 1 mM MgSO_4_, 30 µM H_3_BO_3_, 10 µM MnSO_4_, 1 µM ZnSO_4_, 1 µM CuSO_4_, 0.03 µM (NH_4_)_6_Mo_7_O_24_, 50 µM FeNaEDTA) supplemented with 10 mM MES-Tris pH 5.6 and 0.8 % phytoagar. After six days of stratification in the dark at 4 °C, seedlings were grown vertically in a Conviron CMP 6010 growth chamber (16 h day/8 h night, 22 °C, 65 % relative humidity, photon fluence rate 150 µmol m^-2^ s^-1^). After four days, seedlings were transferred to microscope dishes (MatTek) containing 200 µL polymerized LAK media supplemented with 10 mM MES-Tris pH 5.6 and 0.7 % low melting point (LMP) agarose (Carl Roth). Seedlings were incubated vertically overnight in the growth chamber until the microscopic experiments were conducted.

Before microscopic analyses, seedlings on microscope dishes were placed horizontally and topped with 90 µL imaging buffer (Allen et al., 2001; 5 mM KCl, 50 µM CaCl_2_, 10 mM MES-Tris pH 5.6). Using a 200 µL pipet tip, seedlings were gently attached back to the LAK media-LMP agarose bed and incubated for 10-50 min for recovery until the GEFI fluorescence emission baseline was stable. Microscopic analyses were performed at Leica SP5 II and Leica SP8 confocal laser scanning microscopes using a 10x air objective and photomultiplier tube detectors (Leica Microsystems). Microscope settings were as follows: image format 1024×178 pixels (1536×256 pixels for RW300 experiment); bidirectional scanning at 400 Hz; zoom 0.75 (SP8) or 1 (SP5 II and RW300 experiment at SP8); pinhole 5 AU; line accumulation 2 (SP5 II) and 1 (SP8); line average 1 (SP5 II) and 2 (SP8); offset −0.4 % for blue, cyan, green and yellow emissions and −0.2 % for red emissions; frame rate 6 sec. Laser intensities and gain settings were optimized for each GEFI and kept stable for all experimental replicates. Emission and excitation settings for each GEFI are listed in Supplemental Data Set 1D. After 4 min baseline recording, chemical treatments were performed by dropping 10 µL of 10-fold concentrated stock solutions (Supplemental Data Set 1E) close to the imaged region.

Image processing and analyses were conducted using Fiji (Schindelin et al., 2012). Image processing included background subtraction (2-4), gaussian blur (1), median (1), 32-bit conversion, thresholding of background noise (2-5) and ratio image calculation for ratiometric GEFIs. Normalized fluorescence intensity (ΔF/F_0_) and emission ratio (ΔR/R_0_) analyses, and root tip localized time-dependent vertical response profiles were generated using a custom build Fiji plugin (will be uploaded to github after article acceptance) that utilizes additional plugins, such as VectorGraphics2D-0.13 (https://github.com/eseifert/vectorgraphics2d) and xchart-3.5.2 (https://github.com/knowm/XChart). Fluorescence emissions (F) and emission ratios (R) were measured as average values from each entire processed movie frame and signal changes (ΔF/F_0_ and ΔR/R_0_) were calculated relative to the average value of a 4 min baseline recording as [F(t)-F(baseline)]/F(baseline)] and [R(t)-R(baseline)]/R(baseline)]. Means and SD of experimental replicates were calculated using Excel (Microsoft). For time-dependent vertical response profiles, root tips were detected within each movie frame and regions of interest (ROIs) were drawn to cover the entire x-axis and a defined area above the root tip. Vertical response profiles were calculated from each movie frame ROI as average of all x-axis pixel values within each y-axis pixel line (similar to the Plot Profile command in Fiji) and plotted in a time-dependent manner. Time-dependent signal change vertical response profiles were calculated using the raw response profiles as a basis and applying the signal change formulas to each y-axis pixel line. Average time-dependent vertical response profiles of multiple experimental replicates were generated using the average Z projection command in Fiji.

### Statistical Analysis

For phenotypic analyses presented in Supplemental Figure 5, Box plot- and statistical analyses using pairwise Tukey test comparisons relative to Col-0 wild type were conducted using OriginPro 2018 (OriginLab Corporation).

## Accession Numbers

The Arabidopsis Genome Initiative locus numbers for the genes used in this article are as follows: *ABI1* (AT4G26080), *APA1* (AT1G11910), *AtPEP1* (AT5G64900), *HSP18.2* (AT5G59720), *LTI6b* (AT3G05890), *PYL1* (AT5G46790), *PYR1* (AT4G17870), *VTI11* (AT5G39510).

## Supplemental Data

**Supplemental Data Set 1.** Lists of materials, imaging settings and chemicals used in this work.

**Supplemental Movie 1.** ABA indicator ABA responses in Arabidopsis.

**Supplemental Movie 2.** ABAleonSD1-3L21-P2A-R-GECO1 in response to ABA and IAA.

**Supplemental Movie 3.** PA-17-P2A-ABAleonSD1-3L21 in response to ABA and ATP.

**Supplemental Movie 4.** R-GECO1-GSL-E^2^GFP in response to IAA and ATP.

**Supplemental Movie 5.** R-GECO1-GSL-E^2^GFP in response to glutamate and ATP.

**Supplemental Movie 6.** PA-17-P2A-Grx1-roGFP2 in response to glutamate and H_2_O_2_.

**Supplemental Movie 7.** PA-17-P2A-roGFP2-Orp1 in response to glutamate and H_2_O_2_.

**Supplemental Movie 8.** R-GECO1-P2A-Grx1-roGFP2 in response to H_2_O_2_.

**Supplemental Movie 9.** R-GECO1-P2A-roGFP2-Orp1 in response to H_2_O_2_.

**Supplemental Movie 10.** R-GECO1 and roGFP2-Orp1 in response to ATP and H_2_O_2_.

**Supplemental Movie 11.** R-GECO1-P2A-roGFP2-Orp1 in response to PEP1 and H_2_O_2_.

**Supplemental Movie 12.** R-GECO1 and Grx1-roGFP2 in response to ATP and H_2_O_2_.

**Supplemental Movie 13.** R-GECO1-P2A-Grx1-roGFP2 in response to PEP1 and H_2_O_2_.

**Supplemental Movie 14.** R-GECO1-GSL-E^2^GFP in response to PEP1 and ATP.

**Supplemental Movie 15.** R-GECO1-P2A-Grx1-roGFP2 in response to GSSG.

**Supplemental Movie 16.** R-GECO1-GSL-E^2^GFP in response to GSSG.

**Supplemental Movie Legends**

## ACKNOWLEDGEMENTS

We thank the groups at COS (Heidelberg) for generous access to equipment and GreenGate modules, Dr. Jana Hakenjos for initial help with ABAleon purifications, Dr. Andreas Meyer (University of Bonn) for roGFP2 PCR templates, Dr. Eugenia Russinova (VIB Gent) for providing the AtPEP1 peptide and Dr. Shintaro Munemasa (Okayama University) for helpful discussions. This work was supported by the Deutsche Forschungsgemeinschaft (DFG WA 3768/1-1) to R.W., (DFG AN 1323/1-1) to Z.A. and (DFG KU 931/14-1) to J.K..

## AUTHOR CONTRIBUTIONS

R.W. conceived the project, generated most of the plasmids and transgenic plants, conducted the in vitro characterization of ABAleons, performed all microscopic analyses and wrote the manuscript. P.K. conducted the characterization of ABA indicators in HEK293T cells and revised the manuscript. Z.A. generated and characterized the transgenic lines RW307 and RW308 and revised the manuscript. C.W. developed the GEFI analyzer Fiji plugin and revised the manuscript. G.B. generated the plasmids indicated with GB and conducted preliminary ABA indicator characterizations. K.L. introduced A-17 and generated PA-17 GreenGate modules. J.K. supervised the ABA indicator characterization in HEK293T cells and revised the manuscript. K.S. supervised and hosted the project.

---

**Supplemental Figure 1.**
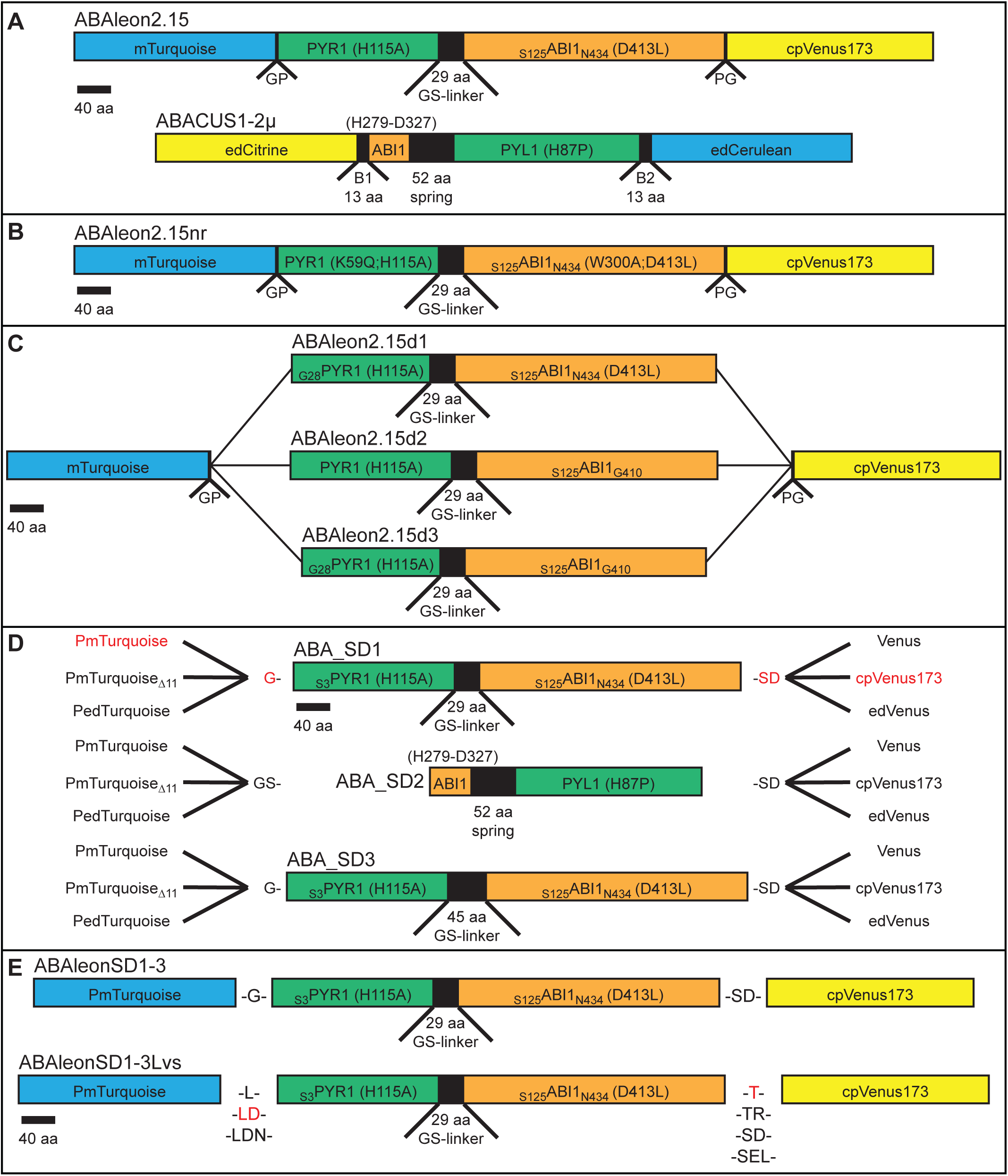
Topologies of ABA indicators. Indicated are fluorescent protein FRET-pairs (cyan and yellow), linkers (black), PYR1 and PYL1 moieties (green) and ABI1 moieties (orange). Point mutations and incorporated amino acids are given. **(A)** ABAleon2.15 and ABACUS1-2µ. **(B)** Non-responsive ABAleon2.15nr that contains two mutations PYR1_K59Q_ and ABI1_W300A_ to prevent ABA-binding. **(C)** ABAleon2.15 deletion variants. **(D)** FRET-pair and sensory domain (SD) variants. Arabidopsis codon-optimized (P)mTurquoise, PmTurquoise_Δ11_ with a C-terminal deletion of 11 amino acids or enhanced dimeric PedTurquoise were used as FRET donor. Venus, circularly permutated Venus (cpVenus173) or enhanced dimeric (ed)Venus were used as FRET acceptor. The sensory domains SD1 and SD3 derived from ABAleon2.15, with SD3 harboring a longer linker between the PYR1 and ABI1 moieties. SD2 derived from ABACUS1-2µ. **(E)** ABAleonSD1-3 linker variants. **(D and E)** Red color indicates the topology of ABAleonSD1-3 **(D)** and the linkers in ABAleonSD1-3L21 **(E)**.

**Supplemental Figure 2.**
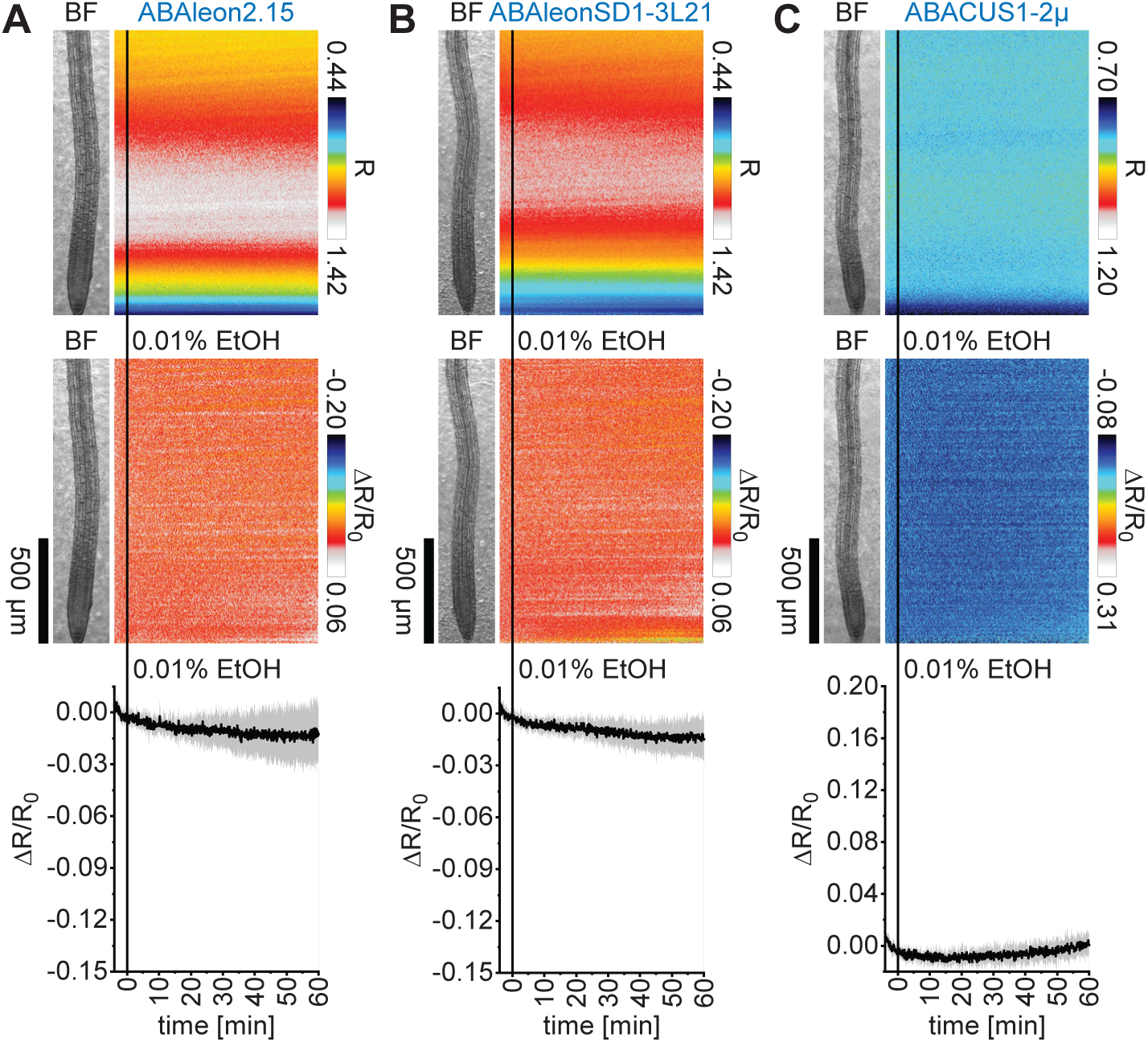
Solvent control experiments of ABA indicators in Arabidopsis. Five-day-old roots of Arabidopsis expressing **(A)** ABAleon2.15 (n = 6), **(B)** ABAleonSD1-3L21 (n = 7) and **(C)** ABACUS1-2µ (n = 6) were imaged for 64 min at a frame rate of 10 min^-1^ and treated with 0.01 % EtOH (solvent control for ABA) at t = 0 min. Shown are average vertical response profiles of (top) emission ratios (R) and (middle) emission ratio changes (ΔR/R_0_) normalized to 4 min average baseline recordings. An adjacent representative bright field (BF) root image is shown for orientation. (bottom) Full image average emission ratio changes (mean ± SD). Color scales of response profiles are identical to the scales in Figure 2.

**Supplemental Figure 3.**
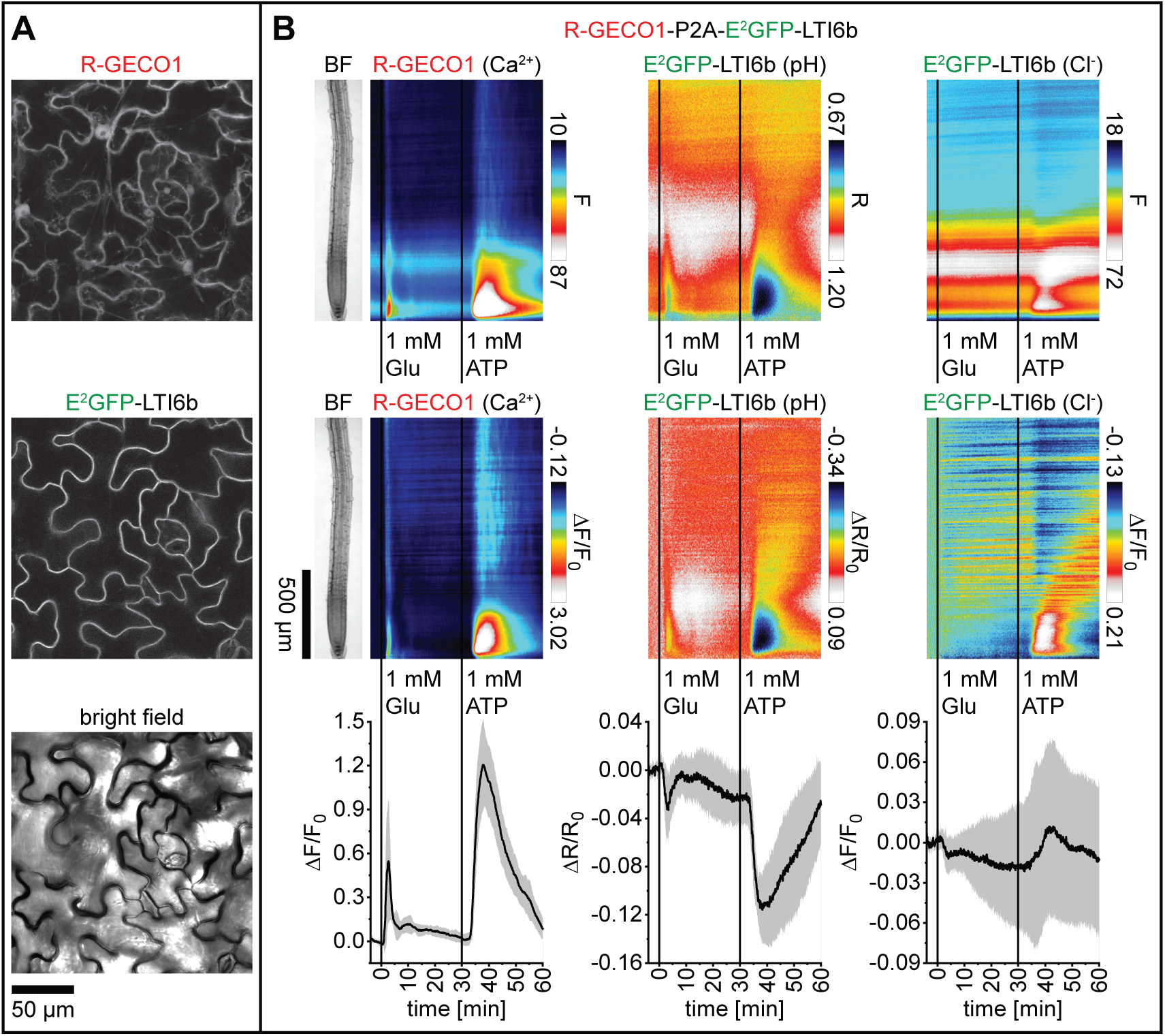
Targeting of E^2^GFP to the plasma membrane. **(A)** Subcellular localization of R-GECO1-P2A-E^2^GFP-LTI6b fluorescence emission in epidermis cells of three-week-old Arabidopsis leaves. **(B)** Analyses of five-day-old roots of Arabidopsis expressing R-GECO1-P2A-E^2^GFP-LTI6b in response to 1 mM glutamate (Glu; t = 0 min) and 1 mM ATP (t = 30 min; n = 8). Images were acquired for 64 min at a frame rate of 10 min^-1^. Average vertical response profiles of (top) fluorescence emissions (F) or emission ratios (R) and (middle) signal changes (ΔF/F_0_ or ΔR/R_0_) normalized to 4 min average baseline recordings. An adjacent representative bright field (BF) root image is shown for orientation. (bottom) Full image signal changes (mean ± SD). These data are related to experiments presented in Figure 5.

**Supplemental Figure 4.**
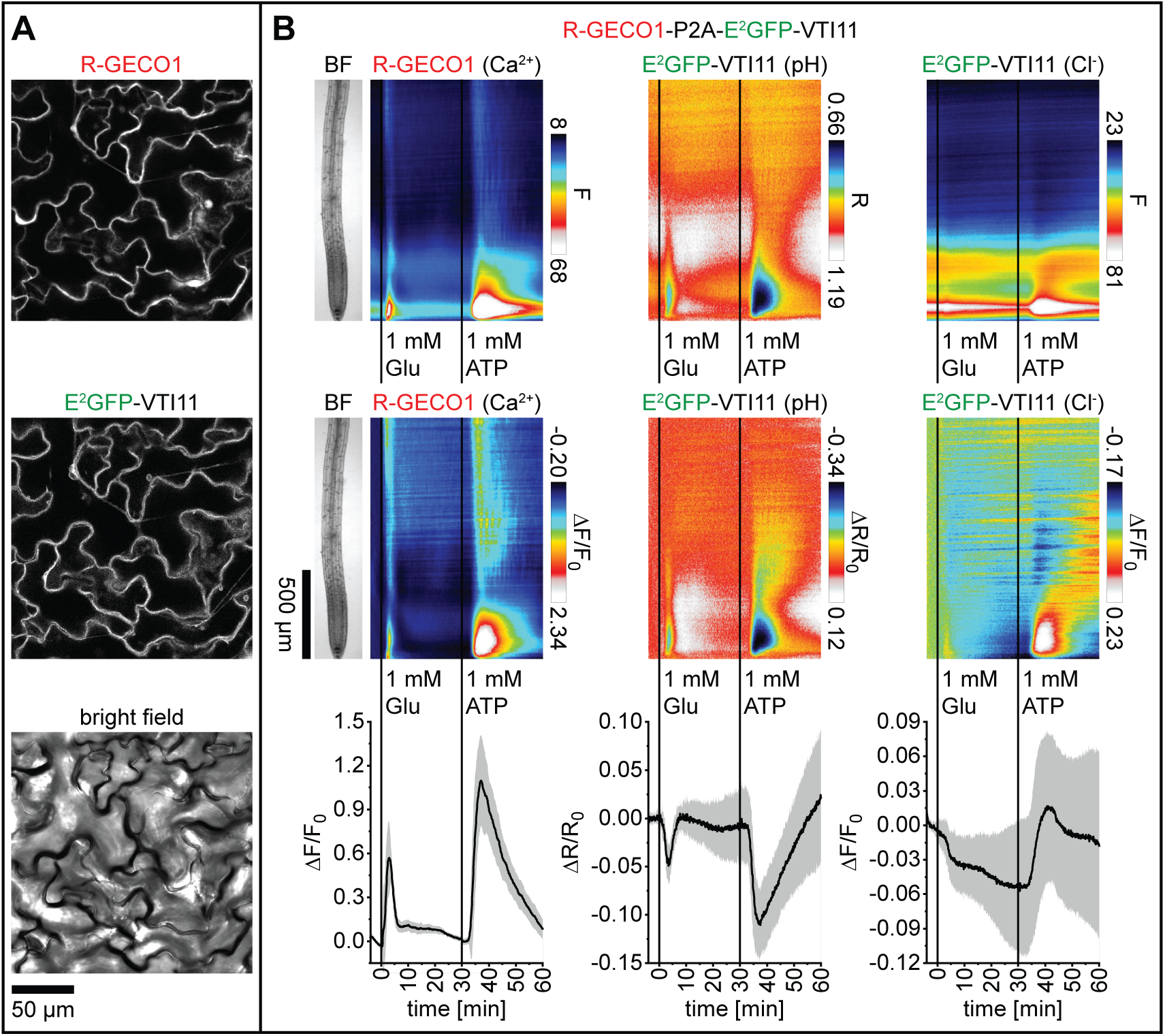
Targeting of E^2^GFP to the tonoplast. **(A)** Subcellular localization of R-GECO1-P2A-E^2^GFP-VTI11 fluorescence emission in epidermis cells of three-week-old Arabidopsis leaves. **(B)** Analyses of five-day-old roots of Arabidopsis expressing R-GECO1-P2A-E^2^GFP-VTI11 in response to 1 mM glutamate (Glu; t = 0 min) and 1 mM ATP (t = 30 min; n = 8). Images were acquired for 64 min at a frame rate of 10 min^-1^. Average vertical response profiles of (top) fluorescence emissions (F) or emission ratios (R) and (middle) signal changes (ΔF/F_0_ or ΔR/R_0_) normalized to 4 min average baseline recordings. An adjacent representative bright field (BF) root image is shown for orientation. (bottom) Full image signal changes (mean ± SD). These data are related to experiments presented in Figure 5.

**Supplemental Figure 5.**
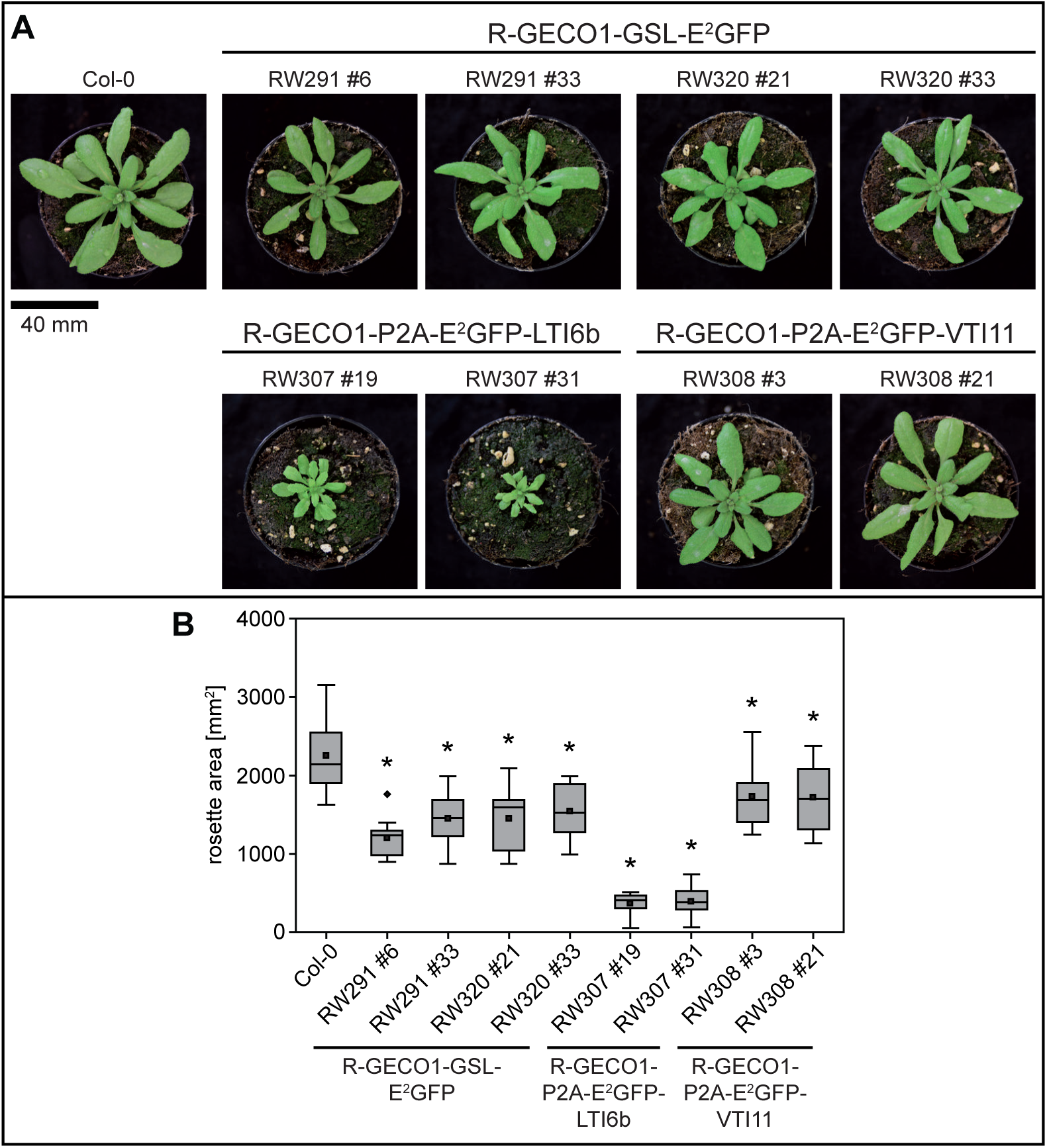
Targeting of E^2^GFP to the plasma membrane induces plant growth defects. **(A)** Representative images of 28-day-old Arabidopsis lines expressing the indicated 2-In-1-GEFIs. **(B)** Rosette area quantification of 28-day-old 2-In-1-GEFI lines (square dots, mean; central lines in boxes, median; boxes, 25^th^ and 75^th^ percentiles, whiskers, ± 1.5 interquartile range; diamond dots, outliers). Asterisks indicate statistically significant differences relative to Col-0 wild type in pairwise Tukey test comparisons (p < 0.05; n = 9-12). These data are related to Figure 5 and Supplemental Figures 3 and 4.

**Supplemental Figure 6.**
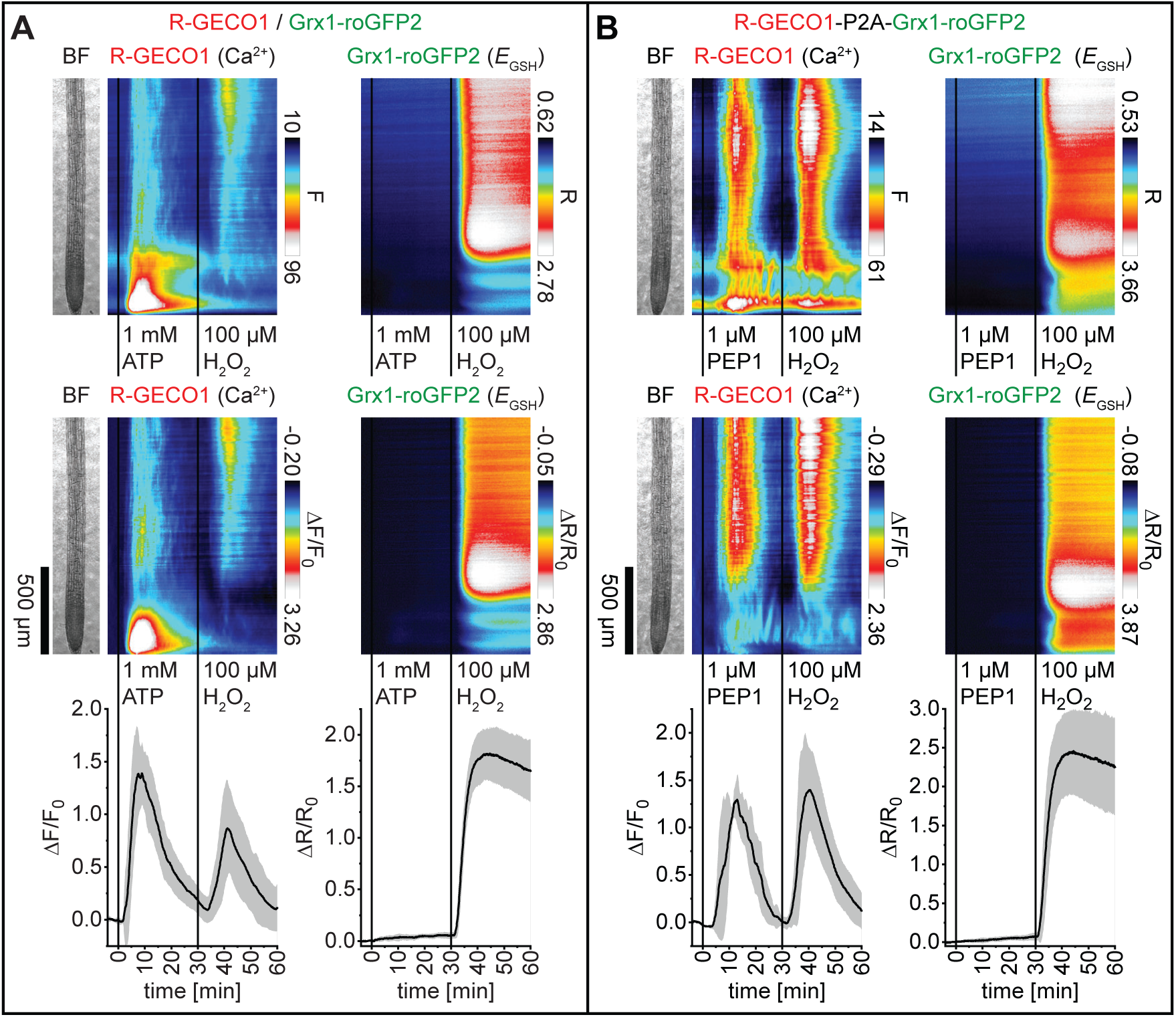
Cytosolic *E*_GSH_ is only weakly affected by ATP and PEP1 treatments. Analyses of five-day-old roots of Arabidopsis expressing **(A)** R-GECO1 and Grx1-roGFP2 (Ca^2+^ and *E* _GSH_; n = 8) in response to 1 mM ATP and 100 µM H_2_ O_2_, and **(B)** R-GECO1-P2A-Grx1-roGFP2 (Ca^2+^ and *E*_GSH_; n = 7) in response to 1 µM PEP1 and 100 µM H_2_O_2_. Average vertical response profiles of (top) fluorescence emissions (F) or emission ratios (R) and (middle) signal changes (ΔF/F_0_ or ΔR/R_0_) normalized to 4 min average baseline recordings. An adjacent representative bright field (BF) root image is shown for orientation. (bottom) Full image signal changes (mean ± SD). Representative experiments are provided as Supplemental Movies 12 and 13. These data are related to experiments presented in Figure 8.

**Supplemental Figure 7.**
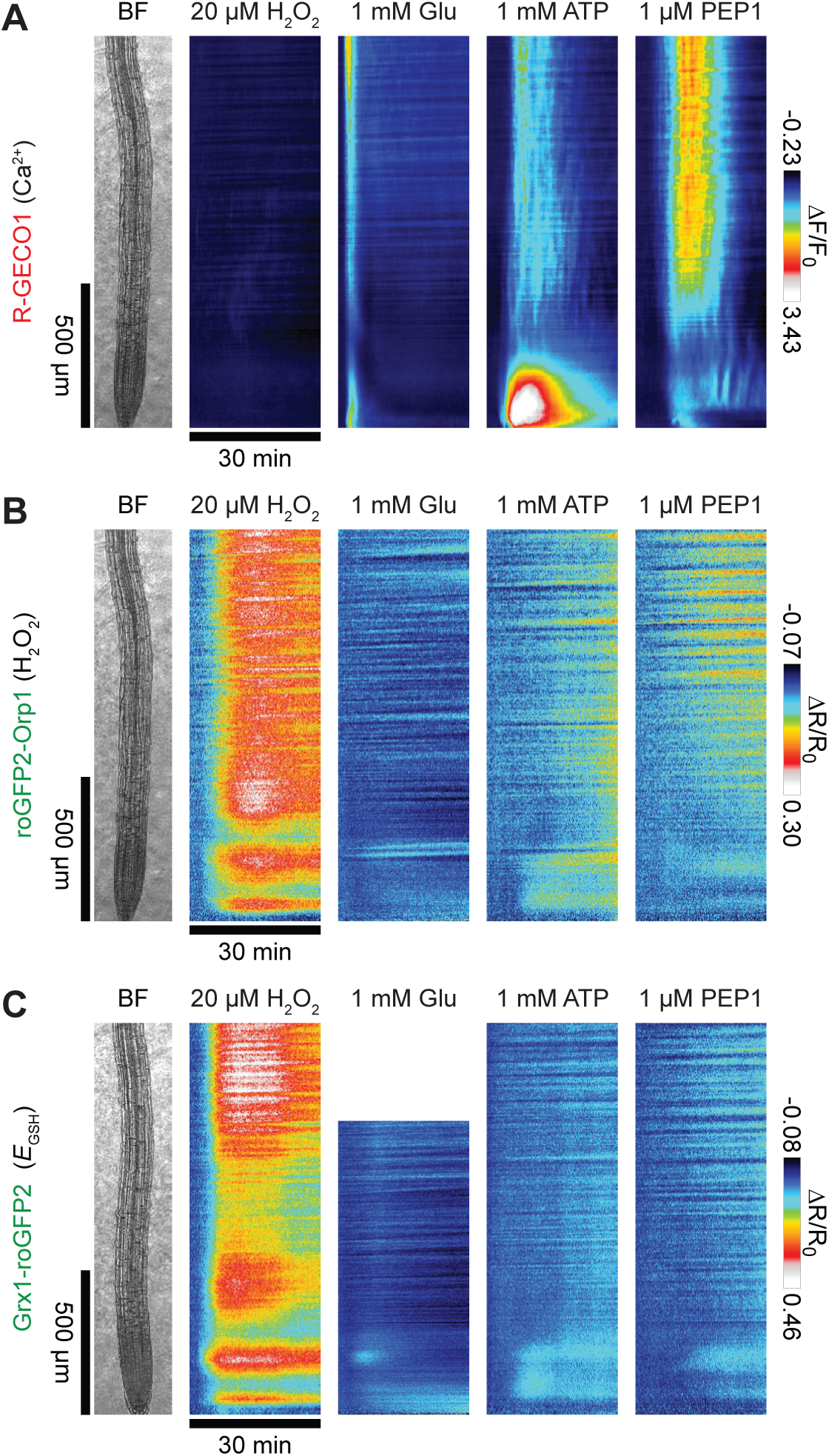
Glutamate-, ATP- and PEP1-dependent cytosolic oxidation is below the threshold of ROS-induced Ca^2+^ signaling. 30 min signal change (ΔF/F_0_ or ΔR/R_0_) (Ca^2+^), **(B)** roGFP2-Orp1 (H O) and **(C)** Grx1-roGFP2 (*E* or ΔR/R_0_) vertical response profiles of **(A)** R-GECO1) (Ca2+), **(B)** roGFP2-Orp1 (H_2_O_2_) and **(C)** Grx1-roGFP2 (*E*_GSH_) after treatments with 20 µM H_2_ O_2_, 1 mM glutamate (Glu), 1 mM ATP and 1 μM PEP1. These data were taken from analyses presented in Figure 7 (H_2_O_2_), Figures 5A and 6B (Glu) and Figure 8 and Supplemental Figure 6 (ATP and PEP1) and scaled to the adjacent color scale.

**Supplemental Figure 8.**
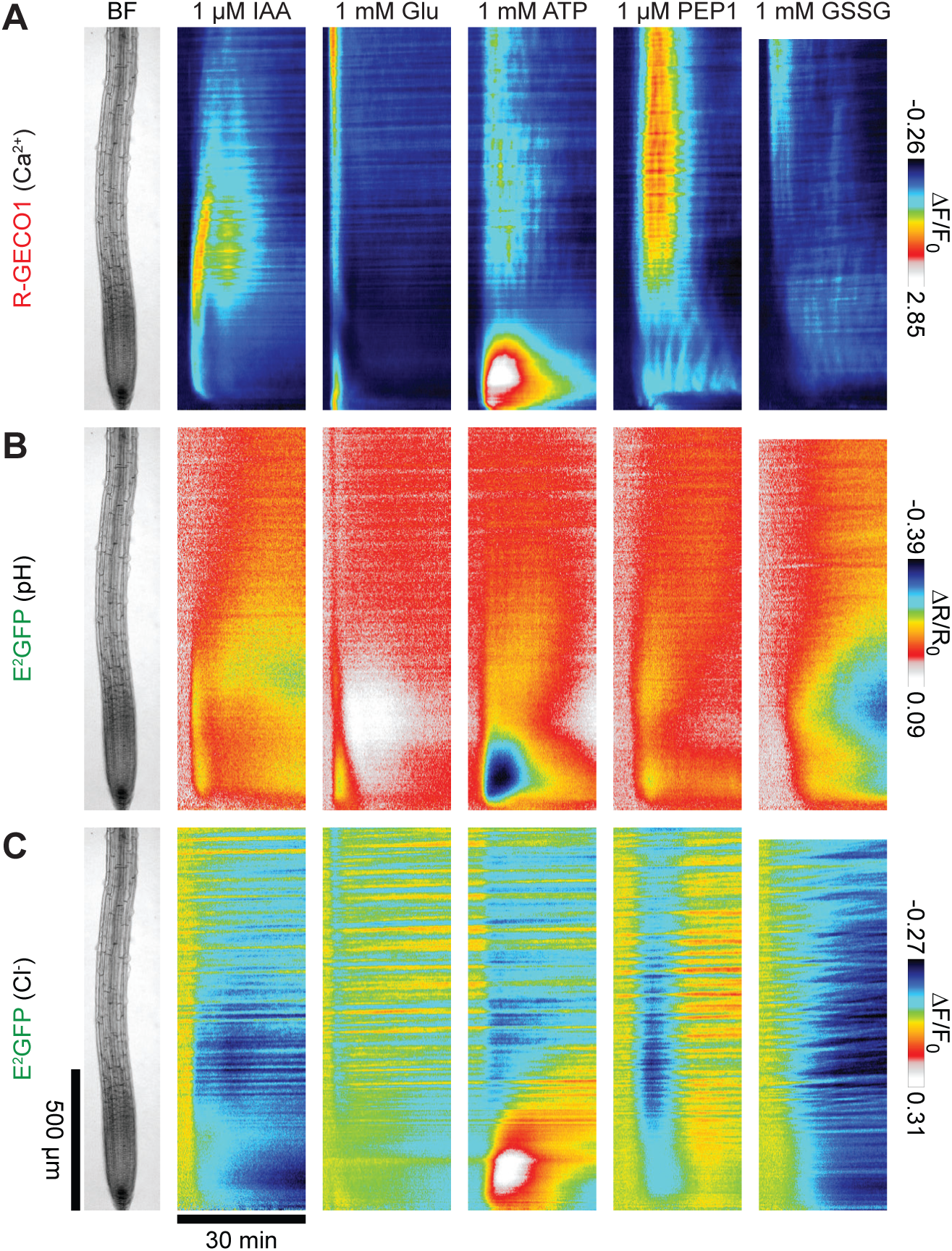
Ca2^+^, H^+^ and anion fluxes exhibit a high spatiotemporal overlap. 30 min signal change (ΔF/F_0_ or ΔR/R_0_) vertical response profiles of **(A)** R-GECO1 (Ca^2+^), **(B)** E^2^ GFP (H ^+^) and **(C)** E^2^ GFP (Cl^−^/anions) after treatments with 1 µM IAA, 1 mM glutamate (Glu), 1 mM ATP, 1 µM PEP1 and 1 mM GSSG. These data were taken from analyses presented in Figure 4 (IAA), Figure 5 (Glu and ATP) Figure 9 (PEP1) and Figure 10 (GSSG) and scaled to the adjacent color scale.

